# Overexpression of MutS impairs DNA mismatch repair and causes cell division defect in *E.coli*

**DOI:** 10.1101/2020.10.13.337683

**Authors:** V Rajesh Iyer, Shivranjani C Moharir, Satish Kumar

## Abstract

MutS and its homologues, from prokaryotes to humans, recognize and bind to DNA mismatches generated during DNA replication, initiate DNA mismatch repair and ensures 100-200 fold increase in replication fidelity. In *E.coli*, through post transcriptional regulation, at least three mechanisms mediate decline of MutS intracellular concentrations during stress conditions. To understand the significance of this multifold regulation, we overexpressed MutS in *E.coli* and found that it led to impairment of DNA mismatch repair as reflected by preferential accumulation of transition mutations in spontaneous base pair substitution spectrum. This phenomenon was dependent on MutS-mismatch affinity and interaction. Higher MutS overexpression levels promoted DNA double strand breaks, inhibited cell division and resultantly caused a manifold increase in *E.coli* cell length. This cell division defect involved a novel MutS-FtsZ interaction and impediment of FtsZ ring function. Our findings may have relevance for cancers where mismatch proteins are known to be overexpressed.

## Introduction

In *E.coli*, mutation is limited to a rate of 10^−9^ to 10^−11^ per generation through the evolution of high fidelity replicative DNA polymerase-III and an efficient mismatch repair system^1^. During replication, a highly conserved, mismatch repair pathway corrects DNA base pair mismatches which are left uncorrected by the proof reading domain of DNA polymerase-III. MutS is a primary component of DNA mismatch repair which recognizes and binds to DNA base pair mismatches and initiates the process of mismatch repair^2^. MutS-mismatch complex enables docking of MutL, which recruits and activates a methylation sensitive endonuclease, MutH, to achieve methyl directed mismatch repair (MMR) in *E.coli*^3,4^. The absence of Dam (DNA-adenine-methyltransferase) mediated methylation on the tetra nucleotide ‘GATC’ of the newly replicated daughter strand, causes MutH to selectively nick the same^5^. Post nicking of daughter strand, through the application of helicase (UvrD), single strand binding protein, exonuclease (RecJ,ExoVII,ExoI or ExoX), DNA polymerase-III and DNA ligase, the entire mismatch repair containing region of DNA is re-synthesized and consequently, error gets corrected^6^.

In *E.coli*, cytokinesis is mediated through a multiprotein complex named FtsZ ring. FtsZ, a GTPase, which upon binding to GTP and through the help of FtsA and ZipA assembles into a ring like structure at the middle of the cell^7,8^. Through GTP hydrolysis, FtsZ enables membrane invagination which subsequently leads to cytokinesis. During cell division, FtsZ ring also acts as a scaffold where proteins required for ring stabilization (FtsQBL, ZapA-D, FtsEX), for chromosome segregation (FtsK), cell wall degradation (envC, amiA, amiB) and septal peptidoglycan synthesis (FtsW) assemble^9^. Under DNA damaging conditions, SOS response induces SulA expression which inhibits FtsZ polymerization^10^, delays cytokinesis and cause cell elongation (a hallmark of SOS response in *E.coli*)^11,12^. Recent reports suggest that DNA double strand breaks can induce SulA independent increase in cell length^13,14^, suggesting, a probable existence of more direct link between DNA damage and FtsZ ring function.

In *E.coli*, intracellular concentration of MutS in stress conditions is regulated through post transcriptional regulation by three mechanisms. First, the stationary phase sigma factor, RpoS, induces expression of small RNA, SdsR, which binds to the coding region of MutS mRNA^15,16^; second, Hfq (RNA chaperone) binds directly to the leader sequence of MutS mRNA^16,17^; and third, Hfq enables small RNA, ArcZ, to bind to the 5’ UTR region of MutS mRNA^17^. Collectively, all three mechanisms in tandem bring down MutS levels under conditions of stress. In order to understand the significance of the three highly evolved, but redundant mechanisms to achieve one goal of MutS reduction in stress conditions, we overexpressed MutS in *E.coli.* We found that MutS-overexpression in *E.coli* leads to specific impairment of MMR as transition mutations dominated the base pair substitution spectrum of spontaneous mutations and this phenomenon was dependent on MutS-mismatch affinity and interaction. MutS-overexpression also led to DNA double strand breaks, caused acute induction of SOS response and induced drastic increase in cell length. Our study also demonstrates that this cell division defect involves a novel MutS-FtsZ interaction and impediment of FtsZ cytokinetic ring function.

## Results

### Overexpression of MutS leads to preferential accumulation of transition mutations

To understand the effect of MutS-overexpression in *E.coli*, we cloned MutS under the influence of a phage derived strong ribosome binding site (sRBS) in an arabinose inducible expression vector (pMutS). Under uninduced conditions, pMutS led to more than 10 folds increase in MutS levels in both exponential as well as stationary phase (Fig. 1A).The overexpression was not toxic to the cells as growth retardation was not observed (Supplementary Fig. 1A). To explore the effect of MutS-overexpression on base pair substitution (BPS) mutations, we conducted Lac-papillation assay^18^, utilizing six strains of *E.coli,each* bearing chromosomally located *lacZ* allele that has a specific loss of function base substitution mutation at 461^st^ codon^19^. The function of these *lacZ*mutant alleles can only be restored through a specific type of base substitution, phenotypically identified by the presence of blue papillae in Lac-papillation assay and their numbers reflect the rate of base substitution. The assay can qualitatively estimate the rate of all the six possible base substitutions, *i.e.*, AT to GC, GC to AT, AT to CG, GC to TA, AT to TA and GC to CG and that, as a whole, conveys total BPS mutation rate. MutS-overexpression led to appearance of a large number of blue papillae specifically in the strain harbouring E461G-*lacZ*-allele which reverts only through GC to AT transition mutation (Fig.1B). The mutation was irreversible as confirmed by the presence of Lac^+^ blue colonies seen upon re-streaking of the blue papillae (Supplementary Fig.1B). The presence of GC to AT transition mutation in the blue colonies was also confirmed by sequencing the E461G-*lacZ*-allele (Supplementary Fig.1C). To validate the above observation, we conducted a fluctuation assay^20^, and observed that upon MutS-overexpression, the growth-dependent GC to AT transition mutation rate was 3 fold higher than control (Fig.1C).

**Figure 1:**
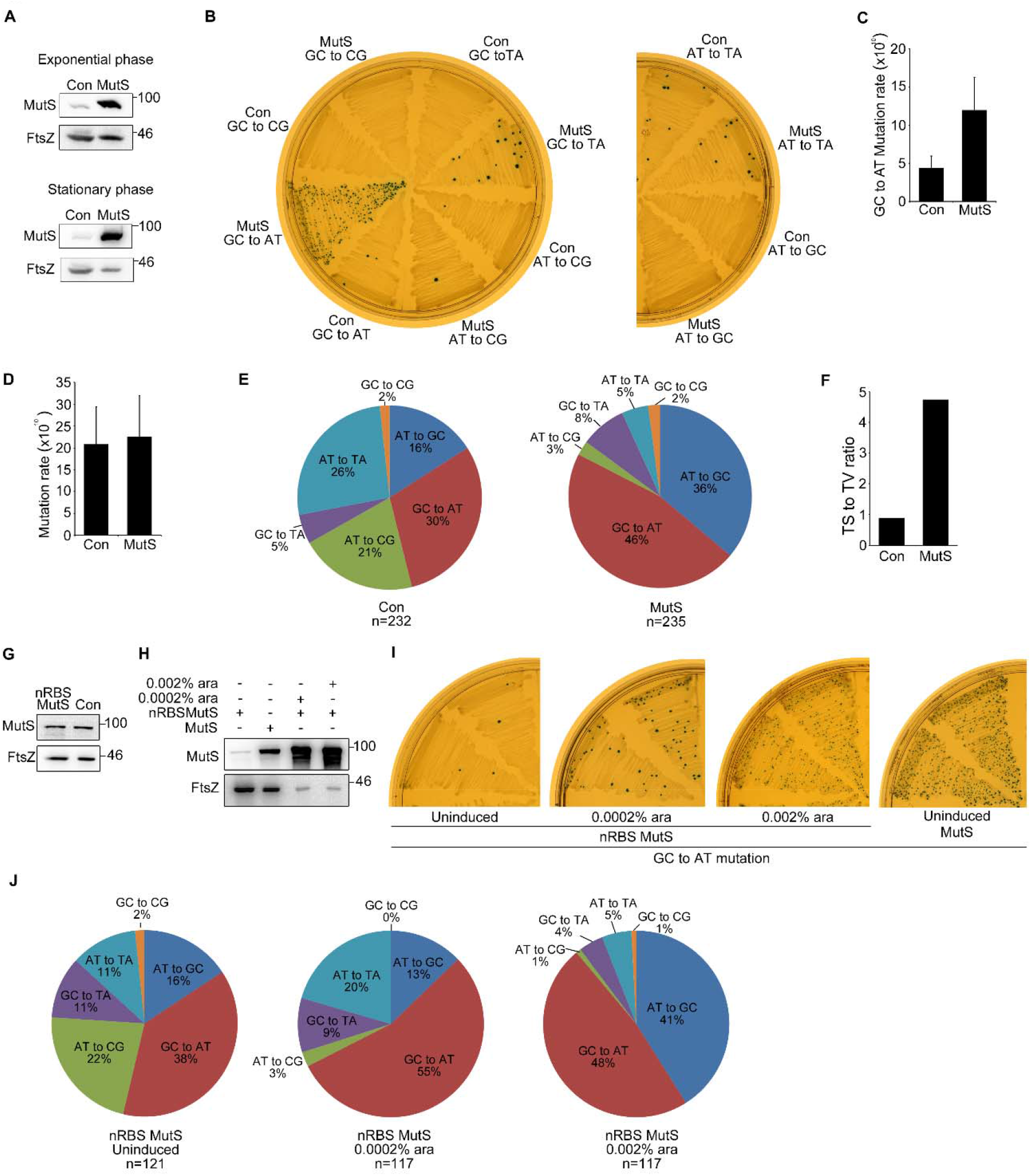
Overexpression of MutS leads to accumulation of transition mutations: **(A)** Western blot showing MutS expression levels in 102BW *E.coli* cells carrying either vector control pBAD18-Kan (Con) or pMutS (MutS) at exponential and stationary phase of growth. **(B)** Representative images of Lac papillation assay, utilizing six *lacZ* alleles to estimate rate of six possible base substitutions, in the presence of either Con or MutS. For each genotype, A single colonywas streaked on LB-Kan Xgal Lactose plates **(C)** Bar diagram showing growth dependent GC to AT mutation rate for E461G-*lacZ*-allele in 102BW cells carrying either Con or MutS plasmids. For mutation rate measurement, fluctuation analysis was conducted with 15 parallel cultures for each group. Error bars represent 95% confidence limits. **(D)** Bar diagram showing mutation rate for spontaneous resistance to rifampicin in 102BW cells carrying either Con or MutS plasmids. For mutation rate measurement, fluctuation analysis was conducted with 19 parallel cultures for each group. Error bars represent 95% confidence limits. **(E)** Pie chart representing base pair substitution spectrum and **(F)** bar diagram representing Ts/Tv ratio of mutations in *rpoB* gene conferring rifampicin resistance in 102BW cells carrying either Con or MutS plasmids. “n” represents the total number of rifampicin resistant clones picked, from five independent experiments for both the groups to be sequenced. (**G and H**) Western blot showing MutS expression levels in exponential phase 102 BW *E.coli* cells with **(G)** vector control (con) or pNRBSMutS (nRBSMutS) and **(H)** pnRBSMutS (nRBSMutS),pMutS (MutS), pnRBSMutS + 0.0002% arabinose, pnRBSMutS + 0.002% arabinose. **(I)** Representative images showing results of *E461G-lacZ-allele* pappillation assay. A single colony of 102BW *E.coli* cells carrying either pNRBSMutS (nRBSMutS) or pMutS (MutS) was streaked on LB-Kan Xgal Lactose plates without (uninduced) or with arabinose (concentration as indicated). Shown are 2 replicates and the number of blue papillae provides an estimate of the GC to AT mutation rate. (J) Pie charts representing base pair substitution spectrum of mutations in *rpoB* gene conferring rifampicin resistance in 102BW cells carrying pNRBSMutS (nRBSMutS), when grown in the absence or presence of the indicated concentrations of arabinose. “n” represents the total number of rifampicin resistant clones sequenced (from two independent experiments for each culture condition). Representative images are selected out of at least three independent experiments.

To probe further, we determined the total base pair substitution (BPS) mutation rate by measuring spontaneous resistance to rifampicin and found no significant change between control and MutS-overexpression (Fig.1D). Since rifampicin resistance (Rif^R^) can occur through 71 possible base substitution mutations in *rpoB* gene^21^, we analyzed the BPS-spectrum leading to it. The BPS-spectrum leading to Rif^R^ differed significantly between control and MutS-overexpression (χ^2^=96.44;*df*=5; P<0.001). We observed that in 82% (194/235) of rifampicin resistant clones selected out of MutS-overexpressed population, mutation leading to Rif^R^ arose due to either a GC to AT or AT to GC transition mutations, which was in contrast to the 46% (107/232) found in the control (Fig.1E,Supplementary Table 1). The Transition to Transversion (Ts/Tv) mutation ratio was found to be 0.86 in case of control but under MutS-overexpression, the ratio jumped more than five folds to 4.73 (Fig.1F). Of the total AT to GC mutations, the 1547^th^ nucleotide position hotspot in *rpoB* gene^22^, accounted for 70% (26/37) in control and 79% (67/85) upon MutS-overexpression. This provides a probable reason for the observed increase in AT to GC mutations in Rif^R^ assay but not in Lac-papillation assaywith E461K *lacZ* allelewhich reverts to Lac^+^ through AT to GC mutation.

To further substantiate our interpretation that MutS-overexpression is the reason for the above observed mutations, we replaced the strong ribosome binding site (sRBS) driving MutS expression in pMutS withthe native ribosome binding site (nRBS) of*mutS* locus. Under uninduced condition, there was no extra chromosomal MutS expression from nRBS-MutS construct (Fig.1G) and *E.coli* cells with E461G-*lacZ*-allele, carrying the construct under uninduced conditions, did not exhibit the extensive blue papillationas observed in the cells carryingpMutS(Fig.1I) and the BPS-spectrum leading to Rif^R^ (Ts/Tv≈1) also differed from that of sRBS construct pMutS (Fig.1J). Next, we utilized arabinose to drive MutS-overexpression further through nRBS-MutS construct and found that induction with either 0.0002% or 0.002% arabinose exceeded the uninduced levels of MutS-overexpression from pMutS (Fig.1H). Correspondingly, upon induction with arabinose, blue papillation in E461G-*lacZ*-allele appeared (Fig.1I) and the BPS-spectrum leading to Rif^R^ (Ts/Tv≈5) phenocopied that of pMutS (Fig. 1J&Supplementary Table 2). These results further strengthen our inference that overexpression of MutS is responsible for the accumulation of transition mutations.

It is well established that transition mutations accumulate predominantly upon inactivation of mismatch repair system^22–24^. We analysed the BPS-spectrum leading to Rif^R^ in *ΔmutS E.coli* cells and also found that transition mutations accounted for 99% (91/92) of the total mutations (Supplementary Table 1). Hence, our finding that transition mutations preferentially accumulate upon overexpression of MutS indicates thatthe mismatch repair system gets impaired upon MutS-overexpression in *E.coli.*

### Overexpression of MutS in mutation-accumulation experiment leads to chromosomal deletions

Mutation-accumulation experiment followed by whole genome sequencing (WGS) provides an opportunity to investigate mutations in a non-selective manner at a whole genome level. To understand the effects of MutS-overexpression globally at the genome level without any selective pressure, we conducted mutation-accumulation experiment. We started with 9 control and 19 MutS overexpressing*E.coli* lines, passaged them for 50 days, corresponding to around 1250 generations per line and performed WGS. Like in Rif^R^ experiment, in mutation-accumulation experiment, the net mutation rate per generation per genome for control (1.68 × 10^−3^) and MutS-overexpression (1.8 × 10^−3^) were similar (Table1). The ratio of non-synonymous vs synonymous mutations is a parameter to gauge selection pressure in mutation-accumulation experiment^24^. This ratio was 2.25 for control and 5.0 upon MutS-overexpression. It did not differ significantly from the expected value of 3.25 (χ^2^=2.3; P=0.13 for control and χ^2^=0.792, P=0.37 for MutS) suggesting selection was minimal during the course of the experiment. We observed that in control, GC to AT mutations were the most frequent (6/18) and transition mutations accounted for 50% of BPS mutations (9/18).These results were similar to those obtained in earlier studies with wildtype *E.coli*^24,25^(Supplementary Table 3). The BPS-spectrum upon MutS-overexpression got altered in the favour of transition mutations, as theyaccounted for 65% (18/28) of total BPS mutations (Supplementary Table 3). The Ts/Tv ratio in control was 1.0 (9/9) whereas upon MutS-overexpression the ratio increased to 1.8 (18/10) (Supplementary Table 3). Although statistically insignificant, the data indicates a trend towards accumulation of transition mutations similar to what we previously observed upon MutS-overexpression in LacZ reversion and Rif^R^ assays.

**Table 1:** Parameters of Mutation accumulation experiment conducted in 102BW *E.coli* overexpressing following proteins

| Strain | No. of point mutations | No. of small indels<sup>a</sup> | No. of large deletions<sup>b</sup> | No. of lines | Generation per line | Total generations | BPSs per line | Indels per line | Mutation rate per nucleotide (X 10<sup>10</sup>) | 95% CL | Mutation rate per genome (X 10<sup>3</sup>) | 95% CL |
| --- | --- | --- | --- | --- | --- | --- | --- | --- | --- | --- | --- | --- |
| Control | 18 | 1 | 0 | 9 | 1254 | 11286 | 2 | 0.11 | 3.63 | 2.23 | 1.68 | 1.03 |
| MutS | 28 | 6 | 8 | 19 | 1226 | 23294 | 1.47 | 0.68 | 3.89 | 1.26 | 1.8 | 0.58 |
| MutL | 22 | 1 | 2 | 9 | 1225 | 11025 | 2.44 | 0.33 | 4.89 | 1.85 | 2.26 | 0.84 |
| MutSMutL | 15 | 0 | 0 | 9 | 1299 | 11691 | 1.66 | 0 | 2.87 | 1.62 | 1.28 | 0.75 |

<sup>a</sup>denotes indels of size ≤4 base pairs
<sup>b</sup>denotes large deletion of size ≥ 10bps
pMutS, pMutL and pMutS-MutL plasmids were used for overexpression of MutS, MutL or MutSMutL in 102BW *E.coli* cells respectively; Control represents empty vector control pBAD18-Kan carrying 102BW cells
CL denotes confidence limit. Standard error of mean of total number of mutations per lines was multiplied with critical values of t-distribution to give the 95% CL<sup>22</sup>

There were fourteen indels upon MutS-overexpression, while just a single instance of small Indels (≤4bps) in case of control (Table 1). Of the fourteen Indels, eight were large deletions (≥10bps) and rest six were small indels. Of the eight large deletions, three were located at the repetitive elements; three represented specific deletions of intergenic sequences found in Glutamine tRNA, Tyrosine tRNA and Alanine tRNA cluster respectively; and out of the last two deletions, one was located in the gene *ydbL* and the other represented the removal of a 46 kb region between *lacZ-rclA*(Supplementary Table 3 and 4). It may be noted that deletion of intergenic sequences in tRNA locus were not a result of an adaptive response to reduce MutS-overexpression, as in lines exhibiting the deletion, MutS overexpression levels were similar to an isogenic strain which did not participate in the mutation-accumulation experiment (Supplementary Fig.2). Collectively,thehigher Ts/Tv ratio observed in mutation-accumulation experiment, enhanced GC to AT mutation rate in LacZ reversion assays and the transition mutations dominated Rif^R^BPS-spectrum, made us to conclude that,overexpression of MutS impairsthe mismatch repair system in *E.coli.* The appearance of deletions in multiple independent MutS overexpressing lines also suggested at a possible MutS-overexpression induced genomic instability.

### MutS-overexpression mediated mutation phenotype is dependent on DNA polymerase-III fidelity

In theory, the accumulation of transition mutations seen upon MutS-overexpression could be either due to impaired repairing of mismatches generated during replication or due to introduction of MutS-overexpression mediated replication-independent de novo errors. Since MutS-overexpression preferentially caused GC to AT transition mutations in LacZ reversion assays, we utilized E461G-*lacZ*-alleleto discriminate between the two above mentioned possibilities. DNA-Polymerase-IIIintroduces random errors during replication and these errors are repaired by the mismatch repair system. If such a replication error is introduced at 1385^th^ nucleotide position of native oriented E461G-*lacZ*-allele, during the replication of leading strand, a G/t mismatch gets created. Whereas, in invert orientation the same G/t mismatch would get created during lagging strand replication (Fig.2A)^26^. In both cases, if the G/t mismatch goes unrepaired, the resultant GC to AT mutation would confer Lac^+^ phenotype. It is shown previously that in *E.coli*, the fidelity of lagging strand replication is more as compared to that of leading strand^26,27^. Thus, during replication, E461G-*lacZ*-allele would bear far more G/t mismatches (correspondingly more chances of being Lac^+^) at 1385^th^ position in native orientation as compared to that in invert orientation. If MutS-overexpression mediated mutation phenotype was due to the impaired repairing of replication errors, we reasoned that it should be dependent on the E461G-*lacZ*-allele’s orientation and consequently on replication fidelity. Hence, we overexpressed MutS in *E.coli* MG1655 strain carrying E461G-*lacZ*-allele in either of the two orientations and found that, compared to native orientation, inversion of E461G-*lacZ*-allele reduced the number of blue papillae in Lac papillation assay (Fig.2B). We further confirmed this by measuring GC to AT transition mutation rate under both orientations of E461G-*lacZ*-allele and found that in invert orientation, mutation rate associated with the allele declined (Fig.2C). This was not due to any reduction in MutS-overexpression levels in strain withE461G *lacZin* invert orientation(Fig.2D). This experiment shows that transition mutations observed upon MutS-overexpression are a result of impaired repairing of mismatches generated during replicationby DNA-Polymerase-IIIrather than being introduced de novo by MutS through some unknown mechanism.

**Figure 2:**
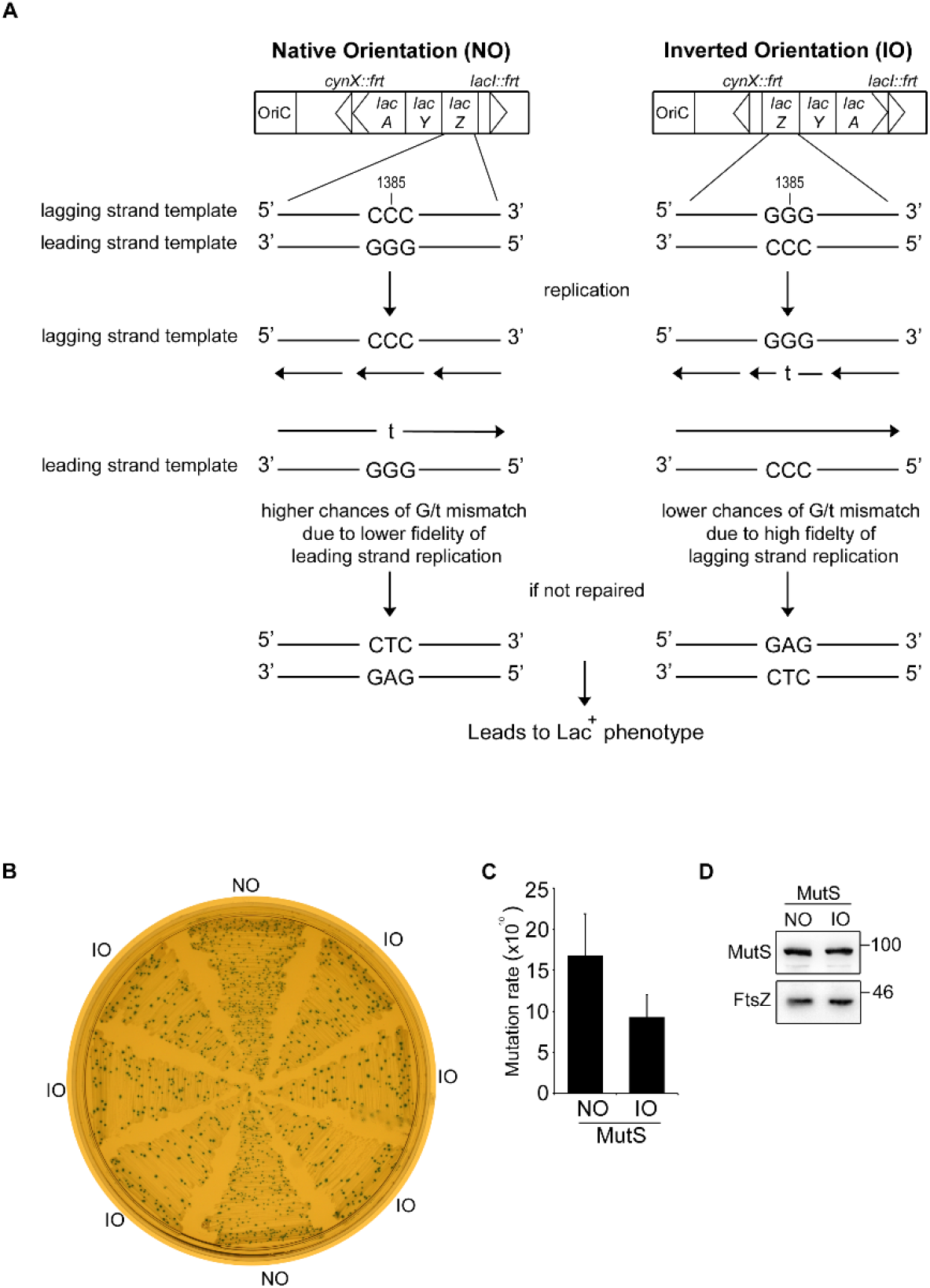
MutS-overexpression mediated mutation phenotype is dependent on DNA-Polymerase-III fidelity. **(A)** Schematic showing introduction of G/t mismatch in E461G-*lacZ*-allele either during leading strand replication in native orientation or lagging strand replication in invert orientation; if unrepaired leads to Lac^+^phenotype. **(B)** Representative image showing result of Lac papillation assay conducted in pMutS carrying MG1655 *E.coli* cells with E461G-*lacZ*-allele in either native orientation (NO, 2 replicates) or invert orientation (IO, 6 replicates). A single colonywas streaked on LB-Kan Xgal Lactose plates. The number of blue papillae provides an estimate of the GC to AT mutation rate. **(C)** Bars shows growth dependent GC to AT mutation rate for either native (NO) or invert oriented (IO) E461G-*lacZ*-allele in MG1655 carrying pMutS. For mutation rate measurement, fluctuation analysis was conducted with 15 parallel cultures for each group. Error bars represent 95% confidence limits. **(D)** Western blot showing MutS expression from pMutS in exponential stage cells of MG1655 *E.coli* cells carrying the E461G-*lacZ*-allele in either native orientation (NO) or invert orientation (IO). Representative images are selected out of at least three independent experiments.

### MutS-overexpression mediated mutation phenotype is dependent on MutS-mismatch affinity and interaction

To probe if MutS-mismatch interaction has any implication on the MutS-overexpression mediated mutation phenotype, we used a mutant of MutS, F36A, which cannot interact with mismatch and is defective in mismatch repair^28^. We overexpressed F36A-MutS in *E.coli* cells carrying E461G-*lacZ*-allele and found that it showed extensively reduced blue papillation as compared to wild type MutS, suggesting that, interaction of MutS with the mismatch is necessary for the accumulation GC to AT transition mutations (Fig.3A). Overexpression of F36A-MutS did not significantly alter Rif^R^ mutation rate (Supplementary Table 5) but BPS-spectrum leading to Rif^R^was significantly different than that of MutS-overexpression (χ^2^=147.80; *df*=5; P<0.001). We found that unlike MutS, F36A-MutS-overexpression did not lead to an accumulation of transition mutations (Fig.3B). Interestingly, in Rif^R^BPS-spectrum of F36A-MutS, transversion mutations (65%, 119/183) exceed transition mutations (35%, 64/183), with AT to TA BPS accounting for 80% (96/119) of total transversion mutations. We utilized another mutant MutS^N^ in which N terminal beta clamp interaction motif is dysfunctional. This mutant is non-functional in mismatch repair^29,30^. We found that overexpression of this mutant MutS^N^ led to almost no blue papillation of cells in E461G-*lacZ*-allele (Fig.3A). Western blots confirmed that the overexpression levels of both mutants, F36A-MutS and MutS^N^, were similar to MutS-overexpression.(Fig.3C). These experiments suggest that MutS-overexpression mediated mutations are dependent upon MutS-mismatch interaction and require MutS functionality.

**Figure 3:**
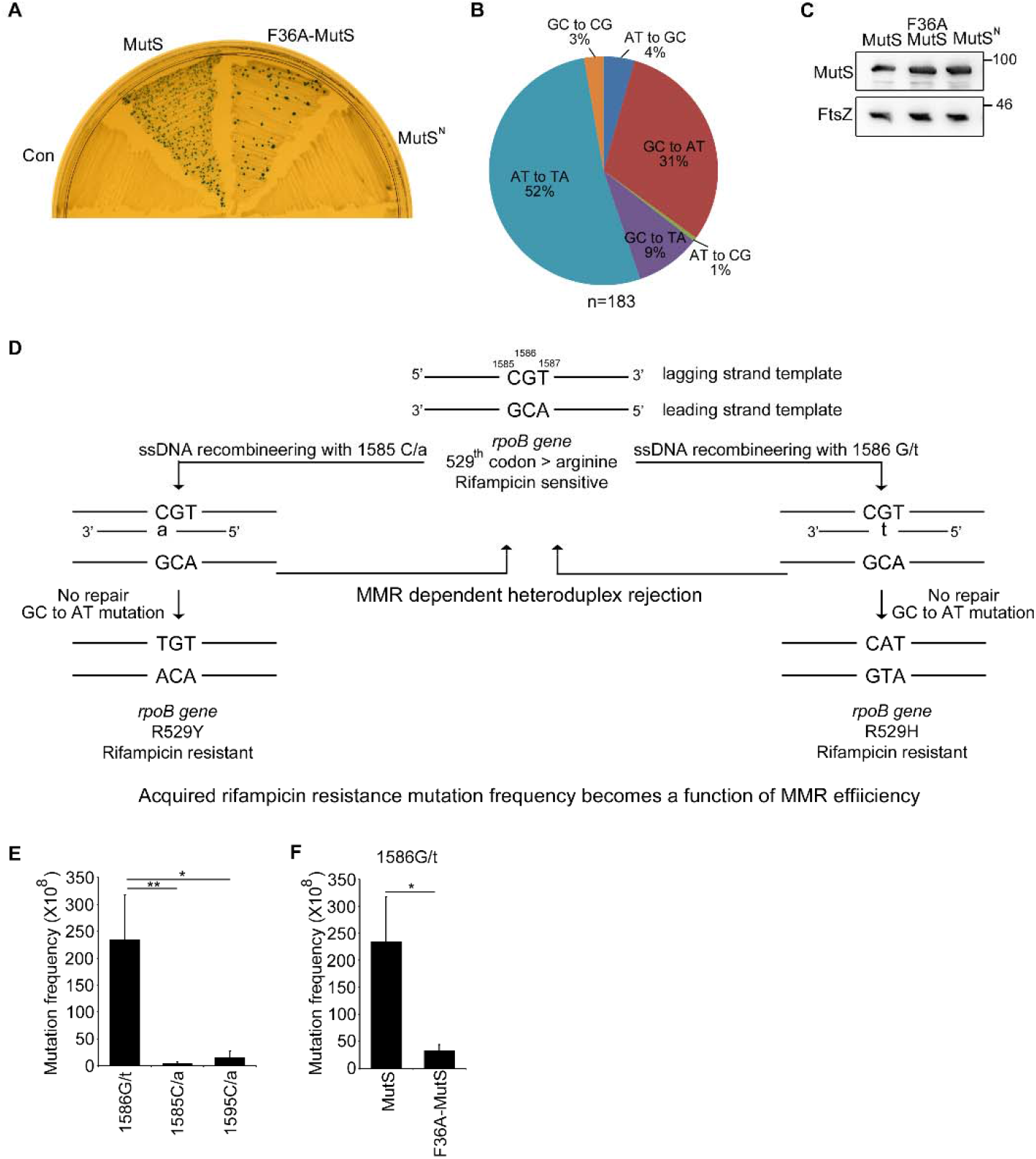
MutS-overexpression mediated mutation phenotype is dependent on MutS-mismatch affinity and interaction. **(A)** Representative images of lac papillation assay in E461G-*lacZ*-allele. A single colony of 102BW cells carrying either control pBAD18Kan(Con),pMutS (MutS),pF36AMutS (F36AMutS) or pMutS^N^ was streaked on LB-Kan Xgal Lactose plates. Plates were incubated for 48 hrs. The number of blue papillae provides an estimate of the GC to AT mutation rate.**(B)** Pie chart represents base pair substitution spectrum of mutations in *rpoB* gene conferring rifampicin resistance in 102BW cells carrying pF36AMutS.“n” represents the total number of rifampicin resistant clones picked, from two independent experiments, to be sequenced. **(C)** Western blot showing MutS expression from exponential phase 102 BW *E.coli* cells carrying pMutS (MutS) or pF36AMutS (F36AMutS) or pMutS^N^ **(D)** Schematic detailing the design of ssDNA oligo recombineering experiment executed to understand the role of MutS-mismatch interaction in MutS overexpression mediated mutation phenotype. Representative images are selected out of at least three independent experiments.**(E)** Bar diagram representing Rif^R^ mutation frequencies observed in pMutS carrying DY378 *E.coli* cells after recombineering with either 1586G/t, 1585C/a or 1595C/a.**(F)** Bar diagram representing Rif^R^ mutation frequencies observed in either pMutS (MutS) or pF36A-MutS (F36A-MutS) carrying DY378 *E.coli* cells after recombineering with 1586G/t.. For **(E)** and **(F),** student’s unpaired two-tailed t-test was used for calculating statistical significance.(*) P < 0.05, (**) P <0.01.

In vitro, MutS has highest affinity towards G/t mismatch^31,32^ and we wondered whether, the MutS-mismatch affinity has any implication in the mutation phenotype. To test this hypothesis, we used ssDNA recombineering to create specific mismatches at *rpoB* gene of *E.coli*^33^, which when not repaired will lead to a transition mutation conferring Rif^R^. We designed two 49 bases long ssDNA oligos which targeted the lagging strand template of *rpoB* gene and upon annealing, created either a C/a mismatch at 1585^th^ (1585C/a) nucleotide or a G/t mismatch at 1586^th^ (1586G/t) nucleotide position of *rpoB* gene respectively (Fig.3D). After recombination, the observed Rif^R^ mutation frequencyprovides a measure through which the mismatch repair efficiencies on these synthetically createdmismatches can be estimated. In control, recombination with 1586G/t lead to seven folds increase in Rif^R^ mutation frequency when compared with that of 1585C/a (Table 2). However, upon MutS-overexpression, recombination with 1586G/t yielded fifty folds higher Rif^R^mutation frequency when compared with that of 1585C/a (Fig. 3E,Table2). We sequenced the *rpoB* gene of these rifampicin resistant colonies and the mutation leading to Rif^R^ was an isogenic GC to AT transition mutation which mapped to the 1586^th^ nucleotide position. Even though Rif^R^ was assayed at 32°C, we thought that the temperature sensitive Rif^R^ conferred by 1585C/a oligo^22^ was a probable cause for the seven folds difference in mutation frequency observed between 1585C/a and 1586G/t in control (Table 2). Thus,we repeated the same experiment with 1595C/a oligo, which upon annealing generates a C/a mismatch at the 1595^th^ nucleotide position of *rpoB* gene and if not repaired, confers a temperature insensitive Rif^R^. In control, compared to 1586G/t,no significant difference in Rif^R^ mutation frequencies was observed upon recombineering with1595C/a. In case of MutS-overexpression, compared to 1586G/t, significant reduction in Rif^R^ mutation frequency was observed upon recombineering with1595C/a (Fig. 3E, Table 2). Next, we conducted recombineering with 1586G/t oligo upon F36A-MutS overexpressionand observed that,compared to MutS, F36A-MutS showed significant decrease in Rif^R^ mutation frequency(Fig.3F, Table 2), thus reconfirming that interaction of MutS with G/t mismatch upon recombineering is necessary for this phenotype. Our results provide evidence for a fifteen folds reduction in efficiency of G/t mismatch repair in a recombination intermediate upon MutS-overexpression. These experiments suggestthat MutS-overexpression can selectively prevent the repair of G/t mismatch for which it poses the highest affinity, and therefore,strongly indicates that MutS-overexpression mediated mutation phenotype is mismatch specific, governed by the affinity of MutS for the mismatch and is not due to any general decline in DNA mismatch repair.

**Table 2:** Outcome of ssDNA recombineering in *rpoB* gene

|  | ssDNA oligo used for recombineering | Phenotype | Rif<sup>R</sup> Mutation frequency (X 10<sup>8</sup>) ± SD* |
| --- | --- | --- | --- |
| **Control** |  |  |  |
|  | 1585C/a | Rif<sup>R</sup> | 3.35 ± 1.76 |
|  | 1586G/t | Rif<sup>R</sup> | 23.44 ± 10.27 |
|  | 1595C/a | Rif<sup>R</sup> | 8.41 ± 7.2 |
| **MutS Overexpression** |  |  |  |
|  | 1585C/a | Rif<sup>R</sup> | 4.17 ± 3.47 |
|  | 1586G/t | Rif<sup>R</sup> | 234.35 ± 83.53 |
|  | 1595C/a | Rif<sup>R</sup> | 15.73 ± 11.78 |
| **F36A-MutS Overexpression** |  |  |  |
|  | 1586G/t | Rif<sup>R</sup> | 32.99 ± 10.87 |

\*Post electrophoration of the ssDNA oligo, cells were grown for 4.5 hrs under non selective conditions and then plated on 50µg/ ml rifampicin containing LB plates and incubated at 32°C. Post 24hrs, colonies were scored. Mutation frequency = count of rifampicin resistant colonies/no.s of viable cells. Shown is the mean ± SD, n=3.
Control represents empty vector control pBAD18-Kan carrying DY378 cells. pMutS and pF36AMutS plasmid were used for overexpression of MutS and F36A MutS in DY378 E.coli cells respectively.

### Co-overexpression of MutL mitigates MutS-overexpression mediated mutation phenotype

During methyl directed mismatch repair, MutS recognizes and binds to the mismatches and enables docking of MutL, which in turn mediates MutH recruitment and activates it to selectively nick the daughter strand^34^. As MutS mismatch interaction was necessary for MutS-overexpression mediated mutation phenotype, we questioned whether MutL and MutH contribute to this phenotype. Through transcriptional fusion, we co-overexpressed either MutL or MutH with MutS as an operon and tested for Lac-papillation in cells carrying E461G-*lacZ*-allele. Interestingly, MutL co-overexpression with MutS completely rescued the mutation phenotype as seen by drastic reduction in blue papillation (Fig.4A) whereas MutH co-expression did not have any effect. Western blots confirmed that this effect was not due to reduction in MutS-overexpression levels upon co-expression of MutL (Fig.4B). Compared to MutS-overexpression, no decrease in growth rate was observed upon MutS-MutL co-overexpression and thus, increased cell death cannot be a cause for the observed rescue (Supplementary Fig.3). When compared to MutS-overexpression, the Rif^R^ mutation rate upon MutS-MutL co-expression remained unchanged (Supplementary Table 5) but the BPS-spectrum leading to Rif^R^ altered significantly (χ^2^=16.11; *df*=5; P=0.006) and exhibited a increase in the percentage of transversion mutations leading to Rif^R^ (18% in MutS, 30% in MutS-MutL) (Fig. 4C)

**Figure 4:**
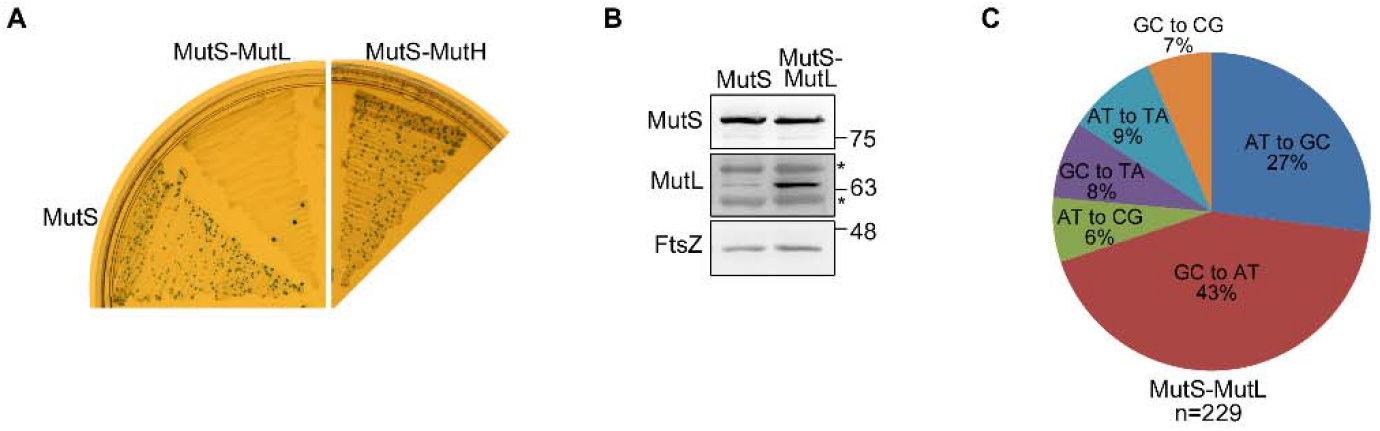
Co-expression of MutL mitigates MutS-overexpression mediated mutation phenotype: **(A)** Representative image of Lac papillation assay in E461G-*lacZ*-allele. A single colony of 102BW cells carrying either pMutS (MutS), pMutSMutL (MutS-MutL) or pMutS-MutH (MutS-MutH)was streaked on LB-Kan Xgal Lactose plates. The number of blue papillae provides an estimate of the GC to AT mutation rate. **(B)** Western blots showing expression of MutS and MutL from exponential phase 102BW *E.coli* cells carrying pMutS or pMutS-MutL. * represents non-specific bands. **(C)** Pie chart represents base pair substitution spectrum of mutations in *rpoB* gene conferring rifampicin resistance in 102BW cells carrying pMutSMutL. “n” represents the total number of rifampicin resistant clones picked, from three independent experiments, to be sequenced. Representative images are selected out of at least three independent experiments.

We also conducted a mutation-accumulation experiment which had 9 lines of *E.coli* cells overexpressing both MutS-MutL and other 9 lines expressing MutL alone. The net mutation rate per genome for MutS-MutL overexpressing group (1.33 × 10^−3^)was slightly lower than that of MutL overexpression [2.26 × 10^−3^] (Table1). In terms of point mutations, MutL overexpressing group behaved similar to vector control group with transition mutation accounting for 54.5% (12/22) of total mutations with GC to AT mutations being the most frequent (Supplementary Table 3). In contrast, MutS-MutL co-overexpression group exhibited a spectrum where transversion mutation accounted for 73.33% (11/15) of total mutations (Supplementary Table 3). The Ts/Tv ratio for MutL overexpression group was 1.2 which (resembled control) whereas MutS-MutL co-overexpression group exhibited a ratio of 0.36 (Supplementary Table 3). The distribution of transition and transversion mutations seen upon MutS-MutL co-overexpression in the mutation-accumulation experiment was significantly different than that of MutS-overexpression (*χ^2^_Yates_*= 4.129; *df*=1; P=0.042).We observed 2 cases of large deletions in MutL overexpression group; but unlike MutS-overexpression lines, where we observed 8 large deletions, MutS-MutL co-overexpression did not show any deletions (Supplementary Table 3 and 4). Overall, above results indicate that MutL co-overexpression mitigates MutS-overexpression mediated mutation phenotype.

### Overexpression of MutS promotes quasi stress induced GC to AT mutations

It is shown that GC to AT mutations preferentially accumulates under stress^35^. Our observation that MutS-overexpression enhances growth-dependent GC to AT mutation rate prompted us to investigate its role in stress induced GC to AT mutation in *E.coli* cells carrying E461G-*lacZ*-allele. We conducted fluctuation assay, in which from each parallel culture of either control or MutS-overexpression group, we plated around 10^8^ cells on minimal lactose plates and exposed them to starvation stress. In 2 days, MutS-overexpression led to appearance of numerous Lac^+^ revertant colonies which were not seen in control (Fig.5A). Post 2^nd^ day, we did not see any increase in number of revertants colonies upon further incubation of minimal lactose plates. Upon re-streaking, revertants displayed stable Lac^+^ phenotype (Supplementary Fig.4A,B) and we also confirmed the GC to AT mutation by sequencing the E461G-*lacZ*-allele (Supplementary Fig.4C).It is known that due to the presence of residual amounts of nutrients on minimal lactose plate, cells undergo division^36^ and we observed that both control and MutS overexpressing cells grew at similar rate (Fig.5B). Since we did not observe such a phenomenon when we plated 10^10^ cells for the earlier GC to AT mutation rate measurement experiment (Fig.1C), we asked whether cell crowding diminishes this phenotype. For testing this, we repeated the same experiment in the presence or absence of *Δlac* scavenger cells, and found that addition of scavenger cells completely diminished the phenotype (Fig.5C). This result suggestthat the Lac^+^ revertants observed in the absence of scavenger cells were indeed due to GC to AT mutations which are happening under selection on the minimal lactose plates.In LacZ reversion assay, growth dependent mutations (mutations which happened before selection) arrive in 2 days and do not fit to a Poisson distribution^20^. The mutations observed upon overexpression of MutS under starvation stress are not growth-dependent mutations as (i) the number of mutations in the fluctuation experiments followed a Poisson distribution [χ^2^= 40.85; df=47; P=0.72] (Fig.5D) (ii) after plating 10^8^ cells, in 2 days, on an average 24 to 36 Lac^+^ colonies appeared upon MutS-overexpression; the earlier estimated growth-dependent mutation rate of 1.2 × 10^−9^ cannot justify the observed number of revertants.Additionally, in LacZreversion assays,stress induced mutations (mutations which happen during selection) are Poisson distributed and their appearance starts at day 2 and the number increases till day 7^36^. In case of MutS-overexpression, though mutations were Poisson distributed, maximum number of Lac^+^ colonies appeared at day 2 and after that there was no substantial increase. As these mutations did not follow the characteristics of either growth-dependent or stress induced mutations, we defined them as quasi stress-induced-mutations.It is shown that stress induced mutagenesis requires recombination or induction of error prone polymerase^37^. To test whether MutS-overexpression mediated quasi stress-induced-mutagenesis is dependent on either, we conducted the experiment in Δ*recA*(recombination deficient) and *ΔdinB* (error prone polymerase IV deficient)*E.coli* cells and found their absence to be nugatory (Fig.5E). In totality, above results indicate that in conditions of starvation stress, MutS-overexpression promotes the appearance of quasi stress-induced-GC to AT mutations.

**Figure 5:**
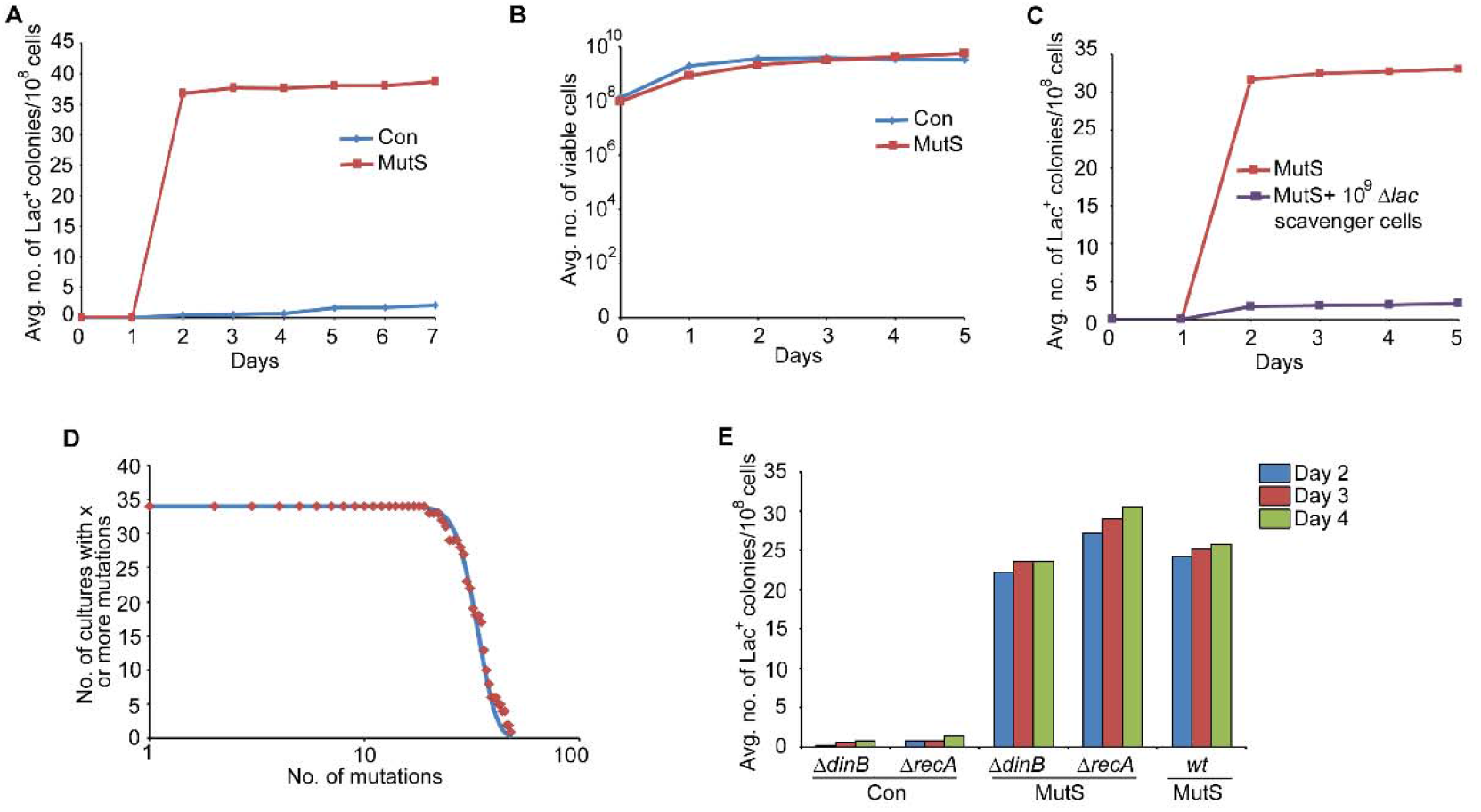
Overexpression of MutS promotes quasi stress induced mutagenesis. **(A)** Plotted data points represent the average number of Lac^+^ revertant colonies counted on 14 minimal lactose agar plates with the passage of days. On Day 0, from 14 parallel cultures of 102BW *E.coli* cells carrying either control pBAD18Kan (Con) or pMutS (MutS), ≈10^8^ cells were spread on minimal lactose agar plates. **(B)** Plotted data points represent average number of viable cells on minimal lactose agar as the days passed. From four minimal lactose plates of either Con or MutS, two agar plugs were taken from each plate at indicated days and suspended and appropriate dilutions were spread on LB agar plates to determine total viable count. **(C)** Plotted data points represent the average number of Lac^+^ revertant colonies observed and counted on 20 minimal lactose agar plates with and without 10^9^ cells of CSH142 (Δ*lac* scavenger cells), with the passage of days. On Day 0, from 20 parallel cultures 102BW *E.coli* cells carrying pMutS, ≈10^8^ cells were spread on minimal lactose agar plates with and without the scavenger cells. **(D)** The figure shows the relationship between number of minimal lactose plates with x or more mutations and log (x). “x” represents the number of Lac^+^ revertants which appeared on minimal lactose agar plates in 2 days from 34 cultures of 102 BW *E.coli* cells carrying pMutS. The line represents the expected number of plates with x or more mutations for a Poisson distribution and the data points represent the actual numbers. **(E)** Plotted data points represent the average number of Lac^+^ revertant colonies counted on 5 minimal lactose plates with the passage of days. On day 0, from 5 cultures of either 102BW *E.coli* cells carrying pMutS (MutS) or VRI5 *E.coli* cells carrying either vector control (Con/Δ*dinB*) or pMutS (MutS/*ΔdinB)* or VRI6 *E.coli* cells carrying either vector control (Con/Δ*recA*) or pMutS (MutS/Δ*recA*), ≈10^8^ cells were spread on minimal lactose agar plates.

### Overexpression of MutS promotes DNA double strand breaks and causes cell division defect

To explore the cause of deletions observed in mutation-accumulation experiment, we considered a possibility of DNA double strand breaks (DSBs) happening in case of MutS-overexpression and tested whether overexpression of MutS enhances susceptibility to radiomimetic agents like bleomycin and zeocin, which intercalates with DNA and cause DNA DSBs^38,39^. Treatment with 0.5μg/ml bleomycin had no effect on both control and MutS overexpressing cells in uninduced conditions (Fig.6A). However, when MutS overexpression levels were elevated using arabinose as inducer(Supplementary Fig.5A), the cells became more susceptible to death at 0.5μg/ml of bleomycin with the maximum effect observed at 0.002% arabinose induction, whereas the control cells remain unaffected (Fig.6B). Similarly, MutS overexpressing cells induced with 0.002% arabinose were alsosusceptible to 0.5μg/ml of zeocin(Fig.6C). It may be noted that DNA DSBs phenotype is sub-optimal under uninduced levels of MutS-overexpression.To validate DNA DSBs occurrence, we overexpressed MutS in strains either predisposed to DNA DSBs (Δ*dam)*or deficient for DNA DSB repair *(ΔruvABC).* In concurrence with the above observations, arabinose induced MutS-overexpression exhibited synthetic sick phenotype with *ΔruvABC* and Δ*dam*mutants of *E.coli* cells (Fig.6D).

**Figure 6:**
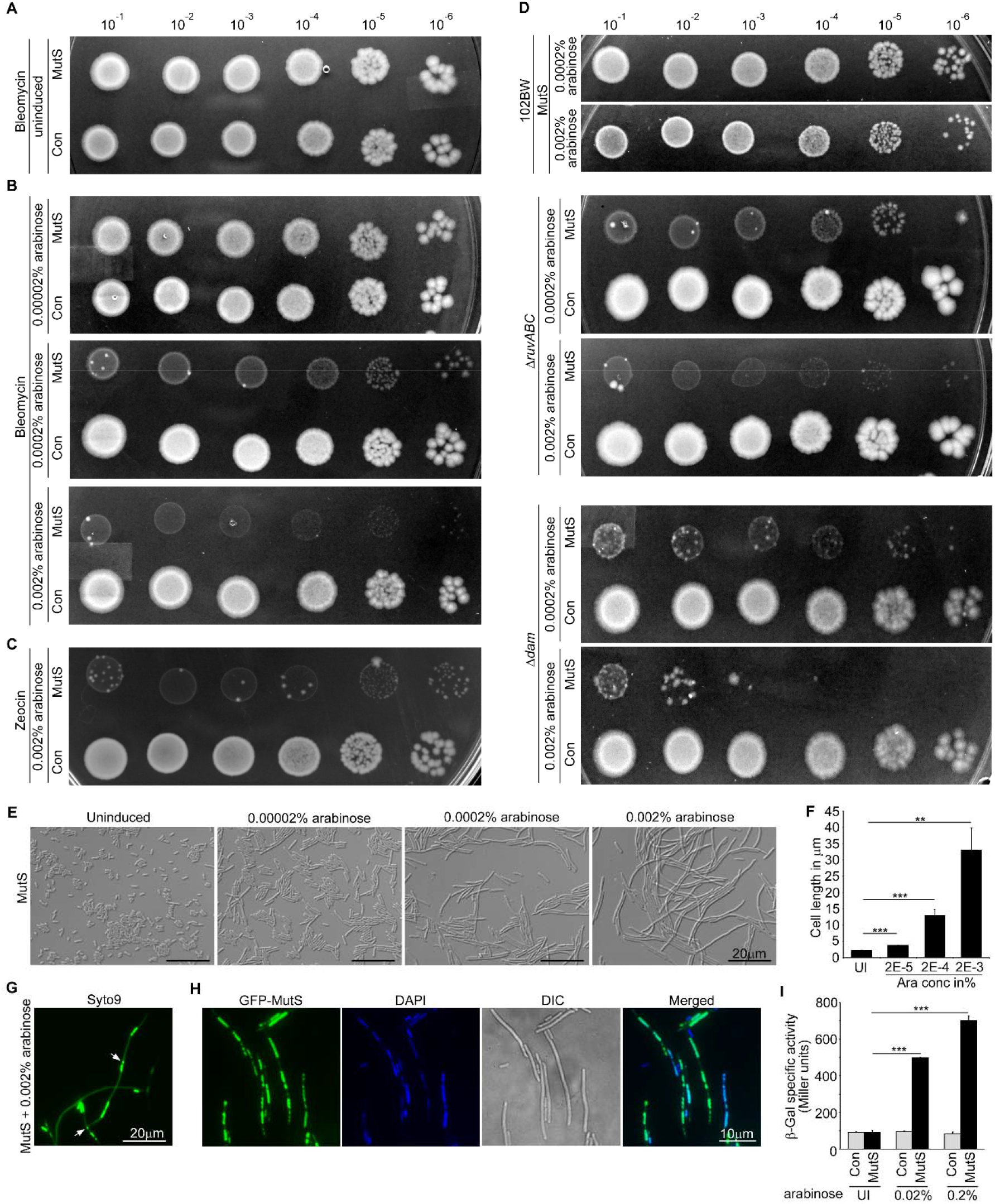
Overexpression of MutS promotes DNA double strand breaks and causes cell division defect. Overnight culture of 102BW *E.coli* cells carrying vector control pBAD18-Kan (Con) or pMutS (MutS) were diluted as indicated and 5 μl of diluted culture was spotted on (A) LB-Kan agar plates with 0.5μg/ml bleomycin, (B) LB-Kan agar plates with 0.5μg/ml bleomycin and indicated concentrations of arabinose, (C) Nutrient Agar-Kan plates with 0.5μg/ml zeocin and 0.002% arabinose. (D) Overnight culture of 102BW, VRI-9 *(ΔruvABC)* and VRI-10 (Δ*dam*) *E.coli* cells carrying either pBAD18Kan (Con) or pMutS (MutS) are diluted as indicated and 5 μl of diluted culture was spotted on LB-Kan agar plates with indicated concentration of arabinose, (E,F,G,H) Overnight culture of *E.coli* cells was diluted 1:1000 in fresh media, shaken till OD_600_≈ 0.1, induced with arabinose (concentration as indicated) and grown till OD_600_≈ 0.6-0.7 (E) Representative DIC microscopy images and (F) cell length analysis of pMutS carrying 102BW *E.coli* cells grown without arabinose (UI) and with arabinose induction at indicated concentrations. For cell length analysis, bars represent mean ± SD; n=3 and 600 cells were counted for each group. Arabninose concentration in % is depicted in scientific notation. (G) Syto9 staining of pMutS carrying 102BW *E.coli* cells grown with 0.002% arabinose. Syto9 (green) stains DNA. Arrows marks indicate fragmented nucleoids (H) Fluorescence and DIC microscopy images of pGFPMutS (GFPMutS) carrying 102BW *E.coli* cells grown with 0.002% arabinose. The cells were fixed and DNA was counterstained with DAPI. GFPMutS is shown as green and DAPI as blue (I) β-galactosidase specific activity of GJ1922 *E.coli* cells *(sulA::lacZ)* harbouring vector control pBAD18-Kan(Con) or pMutS (MutS). Plotted is the average beta galactosidase activity (Miller units) from at least three independent experiments and error bars represent standard deviation. Representative images are selected out of at least three independent experiments. For (F) and (I), student’s unpaired two-tailed t-test was used for calculating statistical significance. (**) P < 0.01, (***) P < 0.001.

Since DSBs induce SOS response^40^ and consequently cell elongation^41^, we examined cell length upon arabinose induced MutS-overexpression. We observed adramatic increase in cell length with increase in concentration of arabinose (Fig.6E,F) which in turn,also led to reduced growth rate (Supplementary Fig.5B). Upon staining DNA with Syto9 stain, each long cell exhibited the presence of multiple nucleoidsindicating inhibition of cytokinesis. We also saw fragmented nucleoids which provide a visual proof for DNA DSBs phenotype (Fig.6G). Under this scenario, in order to understand the localisation of MutS, we overexpressed N-terminal GFP tagged MutS (GFP-MutS) and saw that GFP-MutS localized almost completely on to the nucleoids (Fig.6H). This suggests that MutS binding to DNA contributes to the DNA DSBs and cell elongation phenotype. To confirm whether DNA DSBs induced SOS response, using *sulA::lacZ* fusions^42,43^, we measured SOS response and found that arabinose induced MutS-overexpression elicited significantly higher SOS response than uninduced levels of MutS-overexpression (Fig.6I).

MutS-overexpression mediated cell elongation phenotype had a functional relevance as the mismatch defective mutant, F36A-MutS, failed to show the increased cell length phenotype (Supplementary Fig.5C). To comprehend whether the increased cell length phenotype requires the functions of MutL or MutH, we overexpressed MutS in *ΔmutL* and *ΔmutH* strains and found that the phenotype did not require MutL or MutH functions (Supplementary Fig.5D). To understand the fate of MutS-overexpression mediated mutations in this scenario, we conducted Lac-papillation assay in cells carrying E461G-*lacZ*-allele in the presence of arabinose. At 0.0002% arabinose where the cell length increases substantially, the mutation phenotype disappears (Supplementary Fig.5E) and indicates that cell division defect supersedesthe process of MutS-overexpression induced mutagenesis.In whole, our results suggest that under overexpression, MutS promotes DNA DSBs and cause cell division defect.

### Overexpression of MutS impedes FtsZ ring function

Upon DNA damage, SOS response causes increase in cell length through the expression of FtsZ polymerization inhibitor; SulA^12^. We overexpressed MutS in *ΔsulA* mutant of *E.coli* and found that the cell elongation phenotype was not rescued in *ΔsulA* strain (Supplementary Fig.6A). Cell elongation in *ΔsulA* strains made us to question whether MutS-overexpression can directly influence FtsZ function in *E.coli.* Therefore, we overexpressed MutS in FtsZ-GFP expressing cells and found that in a majority of long cells, either distinct FtsZ ring was not visible or they were carrying multiple FtsZ rings (Fig.7A). To understand, whether this FtsZ ring dysfunction was due to any direct interaction between MutS and FtsZ, we did immuno-precipitation experiments using GFP-MutS and found that MutS interacted with FtsZ (Fig.7B). To understand functional significance of this interaction, we assayed for MutS-overexpression mediated Lac-papillation in E461G-*lacZ*-allelein *E.coli* cells expressing FtsZ-GFP and found it to be completely rescued (Fig.7C). Western blots confirmed that this effect was not due to reduction in MutS-overexpression levels in FtsZ-GFP expressing cells(Supplementary Fig.6B). Additionally, we transformed MutS-overexpressing *E.coli* cells with an origin compatible pTB63 plasmid which expresses the entire FtsQAZ operon and found that it partially alleviated the cell elongation phenotype (Fig.7E,F) and the susceptibility towards bleomycin (Fig.7D). FtsZ ring formation is spatially regulated to preventits formation on the nucleoid^44^. MutS-overexpression did not affect this regulation as we saw that upon counterstaining the DNA, the FtsZ rings still localized to a DNA free region (Fig.7G). The above results indicate that interaction of MutS with FtsZ may impede in FtsZ ring function and consequently, contributes to the mechanism responsible for cell lengthening seen upon MutS-overexpression.

**Figure 7:**
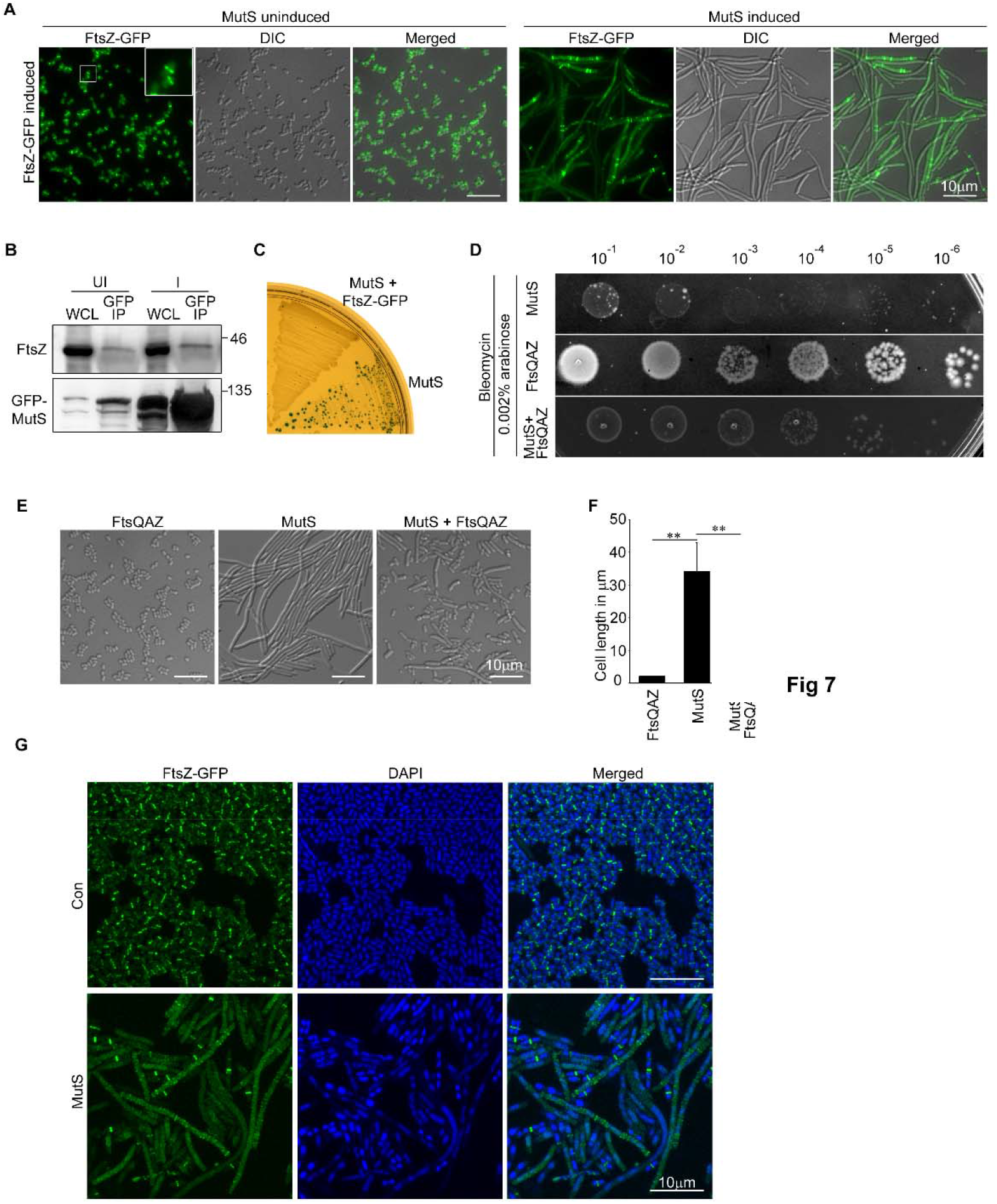
Overexpression of MutS impedes FtsZ ring function. **(A)** Representative fluorescence microscopy images showing FtsZ-GFP rings in VRI8 cells carrying pMutS. Overnight culture of pMutS containing VRI8 *E.coli* cells with chromosomally placed FtsZ-GFP under control of lac promoter, was diluted 1:1000 in fresh media, shaken till OD600≈0.1, induced with either 0.1mM IPTG (MutS uninduced) or 0.1mM IPTG + 0.002% arabinose (MutS Induced), and grown further till OD_600_= 0.6-0.7 **(B)** Western blots showing interaction of FtsZ with GFPMutS. Overnight culture of pGFPMutS containing 102BW *E.coli* cells was diluted 1:1000 in fresh media, shaken till OD600≈0.1, and grown further without (UI) or with 0.002% arabinose induction (I), till OD600= 0.6-0.7. Using GST tagged Anti-GFP nanobody, cell lysates were immunoprecipitated and were analyzed by western blotting. **(C)** Representative image of Lac papillation assay in E461G-*lacZ*-allele. A single colony of pMutS in 102BW (MutS) or pMutS in VRI7 (MutS+FtsZ-GFP) was streaked on LB-Kan Xgal Lactose plates. The number of blue papillae provides an estimate of the GC to AT mutation rate. **(D)** Overnight culture of BW27783 *E.coli* cells carrying either pMutS (MutS), pBAD18-Kan+ pTB63 (FtsQAZ) or pMutS + pTB63 (MutS + FtsQAZ), are diluted as indicated and 5 μl of diluted culture was spotted on LB-Kan agar plates with 0.5μg/ml bleomycin and 0.002% arabinose. (**E)** Representative DIC microscopy images and (**F**) Bars showing cell length of BW27783 *E.coli* cells carrying either pBAD18-Kan+ pTB63 (FtsQAZ), pMutS (MutS) or pMutS + pTB63 (MutS + FtsQAZ). Cells were grown overnight and diluted 1:1000 in fresh media, shaken till OD_600_≈0.1, induced with 0.002% arabinose and grown further till OD_600_= 0.6-0.7. Bars represent mean ± SD; n=3 and at least 600 cells were counted for each group. **(G)** Representative fluorescence microscopy images showing localisation of FtsZ-GFP rings and nucleoids in VRI8 cells carrying either vector control (Con) or pMutS (MutS). Cells were grown and induced as in 7(A). The cells were fixed and DNA was counterstained with DAPI. FtsZ-GFP rings are shown as green and DAPI as blue. Representative images are selected out of at least three independent experiments. For **(F)** student’s unpaired two-tailed t-test was used for calculating statistical significance. (**) P <0.01, (***) P<0.001.

## Discussion

In this study we altered the operational circumstances of a mismatch repair protein MutS, by overexpressing it in *E.coli* and found that it led to impairment of DNA mismatch repair. MutS-overexpression caused an accumulation of transition mutations but the mutation rate, as deduced through Rif^R^ rate and MA experiment, remains unchanged. This can be due to co-occurrence of two events,i) better repair of mismatches leading to transversion mutations. An earlier report suggested that MutS overexpression leads to reduction of GC to TA transversions^45^. Our results extend the same for the rest of transversion mutations except for GC to CG BPS.(ii) Mismatches leading to transition mutations are more prone to MutS-overexpression mediated mutation phenotype. The two phenomena are probably cancelling each other out and hence we did not see any significant increase in mutation rate upon MutS-overexpression.

MutS-overexpression mediated mutations are not a result of denovo errors and their origins lie in the errors committed by DNA-Polymerase-IIIduring replication. The mutations were also not a result of MutS mediated reduced replication fidelity, as overexpression of MutS was sufficient to prevent repair of synthetically created mismatches in the *rpoB* gene. MutS binds to a G/t mismatch with higher affinity than a C/a mismatch^32^ and in recombineering experiments, MutS-overexpression specifically prevented repair of a G/t mismatch. G/t or T/g mismatch, when unrepaired, leads to a transition mutation and we infer that under overexpression, MutS binds to G/t or T/g at a greater frequency than any other mismatch. Thereby increasing the chances of activation of a mechanism by which the repair of G/t or T/g mismatch is hindered and thus we observe the accumulation of transition mutations upon MutS-overexpression.

It is known that MutL prolongs MutS binding to mismatch^46^and is essential for the activation of MutH^47^. If the canonical MMR complex MutS-MutL-MutH were involved in the MutS-overexpression mediated mutation phenotype, co-expression of MutL should have augmented the phenotype, but the observed rescue precludes the involvement of MutL and MutH in the phenotype. We speculate that MutL probably sequesters MutS from activating the mechanism responsible for MutS-overexpression mediated mutation phenotype. Quite similarly, owing to the sequestration of MutS through MutS-FtsZ interaction,cells expressing more FtsZ do not exhibit increased blue papillae in E461G-*lacZ*-allele upon MutS-overexpression.

MutS-overexpression also promoted quasi-stress-induced-GC to AT mutations in E461G*-lacZ-*allele. We speculate that during non-selective growth, MutS-overexpression may prime a very small fraction of cells in a population to undergo GC to AT mutation in E461G-*lacZ*-allele which later under specific conditions, manifest themselves as Poisson distributed quasi stress-induced-mutations. A similar kind of mechanism has recently been proposed for the stress induced mutagenesis in FC40 strain of *E.coli*^48^.

Higher levels of MutS-overexpression in *E.coli* cells led to the generation of DNA double strand breaks, caused an acute induction of SOS response and cell division defect. Recently, it is shown that *E.coli* cells experiencing DNA DSBs exhibit increase in cell length and this is not dependent on the action of SOS induced FtsZ inhibitor SulA^13,14^. Our study provides further evidence for a SulA independent mechanism for cell length increase as MutS overexpression directly impeded with FtsZ ring function. We also uncovered a novel interaction between MutS and FtsZ, which probably contributes to the observed cell division defect. Like many studies where FtsZ ring function is inhibited^44,49,50^, co-expression of FtsQ, FtsA and FtsZ partially rescued the cell length phenotype. How MutS-overexpression impedes FtsZ ring? Towards this question, we can only speculate that, MutS-overexpression prevents maturation of FtsZ ring by preventing recruitment of some essential factors required for cytokinesis orMutS-FtsZ interaction, like SulA-FtsZ interaction,inhibits FtsZ-GTPase activity.

We found two distinct phenotypes upon MutS-overexpression i.e., mutation phenotype and cell elongation phenotype. In yeast, overexpression of the primary mismatch repair complex (Msh2-Msh6) is shown to be mutagenic and the phenotype is attributed to replication fork instability due to enhanced Msh6-PCNA interaction^51^. In *E.coli*, if MutS-overexpression mediated mutations were due to replication fork instability, in our ssDNA recombineering experiments, MutS-overexpression should have equally rejected either G/t or C/a mismatch generating ssDNA oligo. F36A-MutS poses a functional β-clamp interaction motif and yet its overexpression fails to phenocopy MutS-overexpression mediated mutation phenotype.This makes us to incline towards a mechanism where MutS-mismatch affinity governs the appearance of the phenotype.

Based on our observations, we interpret that MutS-overexpression mediated mutation phenotype is the basis for cell length phenotype. Under overexpression, a small fraction of MutS possibly interacts with an endonuclease and the resultant MutS-endonucleasecomplex binds on to a mismatch and the endonuclease nicks the DNA without any regard to parental or daughter strand and resultantly causes mutations. Due to higher affinity of MutS for G/t or T/g mismatches, MutS-endonuclease complex docks on to these mismatches at a higher frequency and resultantly we see transition mutation upon MutS overexpression.Under higher MutS-overexpression levels, the same endonuclease probably leads to DNA DSBs and in conjunction with MutS-FtsZ interaction mediated FtsZ ring inhibition,causes the cell division defect (Fig.8). This also explains why the mutation and cell elongation phenotype seen upon MutS overexpression are mutually exclusive. In mutation-accumulation experiments, we also found three evidences of specific deletion of intergenic sequences between tRNA genes in three separate MutS overexpressing lines and this provides a precursory evidence for a sequence specific endonuclease in the above mechanistic scheme.

**Figure 8:**
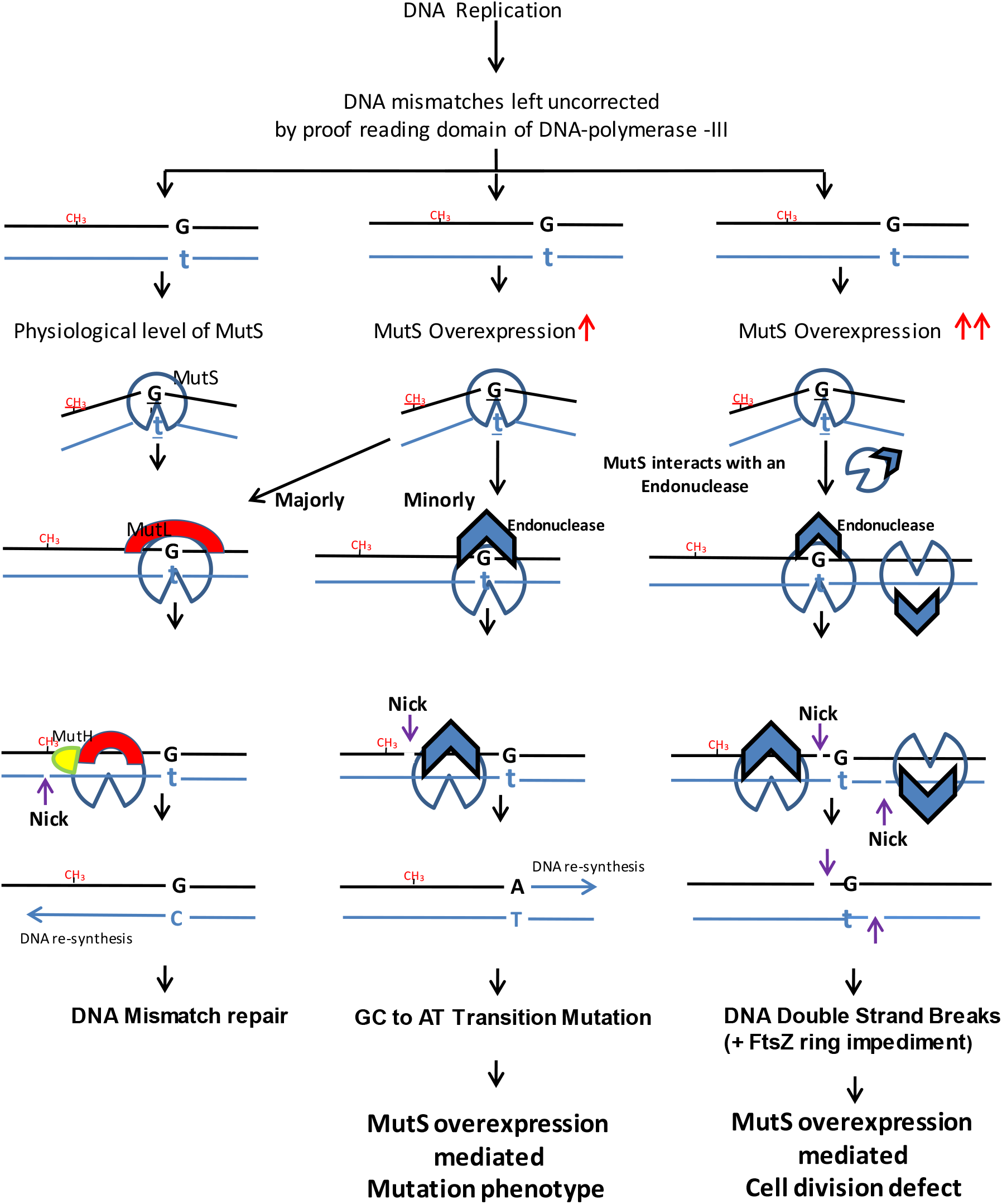
Model depicting the mechanism for MutS overexpression mediated phenotypes in *E.coli.*

We began this study to understand the significance of three mechanisms evolved to regulate MutS expression levels in *E.coli.* Our study implies that these mechanisms may have got evolved to counter the effects of MutS-overexpression observed in this study. Our results are probably amplified versions of phenotypes which may happen in an extremely tiny fraction of *E.coli* cells and contribute to spontaneous mutations. It is well established that, in *E.coli*, spontaneous rate of GC to AT mutations is highest among other substitutions^24^ and we wonder whether it is due to the superfluous MutS presencein a small minority of cells in the *E.coli* population. As MutS overproduction can inhibit cytokinesis, MutS-overexpression may have an effect on generation of antibiotic tolerant persistor cells population^52^.

In many cancers, mismatch repair proteins are overexpressed^53–55^. In Prostate cancer, there are studies which indicate that Msh6overexpression contributes to aggressiveness of the cancer^54,56^. Additionally, mutational signatures in prostate cancer are dominated by GC to AT transition mutations^57^. In eukaryotic mismatch repair, MutS-alpha is a hetero-dimer of Msh2 and Msh6, where Msh6 is the subunit which interacts and binds tomismatch and hence functionally can be compared to MutS in *E.coli*^58,59^. Based on MutS-overexpression mediated transition mutations in *E.coli*, we contemplate that Msh6 overexpression in prostate cancer may be considered as one of the reasons for the appearance of transition mutations dominated mutational signature.

## Materials and Methods

### Strains, molecular biology and media

The *E.coli* strains and plasmids used in this study are mentioned in Supplementary Table 6. The sequence of the primers used for construction of plasmids and recombineering are as mentioned in Supplementary Table 7. The parent strain for most of our experiments was BW27883, which allows homogenous expression of proteins from arabinose inducible pBAD expression vectors^60^. Standardized methods for molecular cloning were utilized during this study. Unless specified, *E.coli* cells were propagated in Difco’s LB broth (Miller) or LB agar (Miller). Nutrient Agar (Himedia laboratories) was used for spot viability assay against Zeocin. Bacto Agar was used at 2% wherever needed. Antibiotics were added as follows: Kanamycin (40μg/ml), Chloramphenicol (10μg/ml), Tetracycline (10μg/ml), Ampicillin (25μg/ml) and rifampicin (50μg/ml). Zeocin, Bleomycin, L-Arabinose and IPTG concentrations are used as indicated. Unless indicated,the cultures were grown at 37°C and shaken at 200-220 rpm.

### Western Blotting

*E.coli* cell lysates were subjected to SDS-PAGE and electro transferred to a 0.22μ PVDF membrane. The membrane was washed with TBST (Tris buffered saline + 0.1% Tween 20) and blocked with 5% blotto for an hour at RT. Anti MutS (US Biologicals M9557-01), Anti MutL (Biorbyt,Orb51498), Anti FtsZ (Biorbyt,Orb 400616), Anti-GFP antibody (Santacruz, sc-8334) were diluted in 1% blotto at 1:3000, 1:3000, 1:5000 and 1:1000 dilutions respectively and membrane was probed overnight at 4°C. Post washing, membrane was probed with HRP conjugated Anti rabbit IgG (Santacruz, sc-2004) at 1:5000 dilution and blot was developed using chemiluminscence. (Santacruz ECL substrate sc-2048; Vilber chemiluminescence imager)

### β-galactosidase assay

β-galactosidase assaywas conducted as described previously^61^. As GJ 1922 cells can utilize arabinose as carbon source, during the experiment, MutS-overexpression levels in these cells were induced with higher concentrations of arabinose.

### Spot Viability assay

Overnight cultures were serially diluted and 5μl of each dilution was spotted on LB agar medium, supplemented as indicated.

### Lac papillation assay

Cells were streaked on LB agar with 0.1% Lactose (Sigma L-3625), 40μg/ml X-Gal (Sigma B4252) and 40μg/ml kanamycin. Plates were incubated in 37°C for 48 hours and photographed.

### Fluctuation assay

For fluctuation assays, overnight grown cells were diluted and roughly cells in thousands were seeded on to multiple parallel cultures (ranged 10-20 culture). The cells were grown at 37°C at 220 rpm. In case of *lacZ* allele, entire culture (10ml) was pelleted and washed twice with 1X M9 salts, plated on M9-lactose plates (M9 media with 0.1% lactose), incubated at 37°C and scored after 48 hours. In case of *rpoB*, 1ml culture was pelleted and spread on LB agar with 50μg/ml rifampicin, incubated at 32°C and scored after 24 hours. Cells were also plated on LB agar to determine viable cell count. Mutation rates and confidence intervals were calculated using the webtool bz-rates^62^.

### Rifampicin resistance spectrum analysis

Colonies which appear in 24 hours on LB rifampicin plates were marked and then plates were kept for additional incubation at 32°C for additional 12 hours. The marked colonies were suspended in 100μl of lysis buffer (20mM Tris, 1mM EDTA, 0.1% SDS, 400mM Nacl, 20μg/ml Proteinase K, 100μg/ml RNaseA, pH 8.0) and kept at 55°C for overnight. The following day, DNA precipitation was carried out by adding 500μl of absolute ethanol and pelleted. The pellet was stripped of any residual ethanol, air dried and dissolved in TE buffer. As according to previous report^22^,colonies were always picked starting from the centre of plate and without any bias for size. Mutations leading to rifampicin resistance in *rpoB* gene fell on to two clusters i.e. cluster I and II and of the two,cluster-II accounts for 90% of mutations^21,22^. Therefore, a primer pair rpoBF and rpoBR was used to amplify the *rpoB* gene region encompassing cluster-II. The PCR product was cleaned up and sequenced. We omitted cluster I as we were able to map more than 99% of mutations to cluster-II. Using MEGA multiple sequence alignmenttool^63^, the sequences were aligned and Rif^R^BPS-spectrum was deduced.

### Construction of plasmids

#### pMutS

Using the primers EcoMutSF and EcomutSR, *mutSgene* was amplified using *E.coli* K-12 MG1655 gDNA and cloned at Nhe-I and Sal-I sites of pBAD18Kan^64^ vector to yield pMutS. Primer “EcoMutSF” contain the strong ribosome binding site (RBS) derived from phage T7 major capsid protein. This RBS is commonly found in all pET vectors. The entire MutS gene was sequenced completely and it was devoid of any mutation.

#### pF36AMutS

Two silent mutation c.37A>G and c.38G>C in were engineered to generate a SacI site in *mutS* gene. This *mutS* gene was cloned at NheI and SalI of pBAD-18Kan to generate psilent-MutS. Through complementation assays in Δ*mutS* mutant of *E.coli*, MutS functionality was validated. Primers EcoMutSF and F36AR were used to introduce F36A mutation and the corresponding amplicon was ligated at Nhe-I and Sac-I sites of psilent-MutS. The F36A-MutS construct was sequenced and verified.

#### pMutS^N^

A 457 bp amplicon was amplified using the primers MutSUPF and phosphorylated MutSNR. This amplicon corresponds to upstream region and first 19 codons of beta clamp mutant *mutS* gene. A 2.5 Kb fragment (rest of *mutS*gene) using phosphrylated MutSNF and EcoMutSR was amplified. These two fragments were ligated and resultant *mutS^N^* gene was re-amplified using primers EcoMutSF & EcoMutSR and cloned at Nhe-I and Sal-I sites of pBAD18kan.The MutS^N^construct was sequenced and verified.

#### pMutL

Using the primers EcoMutLF and EcomutLR, *mutL* gene was amplified using *E.coli* K-12 MG1655 gDNA and cloned at Nhe-I and Sal-I sites of pBAD18Kan vector to yield pMutL. Primer “EcoMutLF” contain the strong ribosome binding site (RBS) derived from phage T7 major capsid protein. The construct was verified by sequencing.

#### pMutS-MutL

Using the primers EcoMutLF-SalI and EcoMutLR, *mutL* gene was amplified using *E.coli* K-12 MG1655 gDNA and cloned at Sal-I site of pMutS vector to yield pMutS-MutL. Primer “EcoMutLF-SalI” contain the strong ribosome binding site (RBS) derived from phage T7 major capsid protein. The construct was verified by sequencing.

#### pMutS-MutH

Using the primers EcoMutHI and EcomutHR, *mutH* gene was amplified using *E.coli* K-12 MG1655 gDNA and cloned at Sal-I site of pMutS vector to yield pMutS-MutH. Primer “EcoMutHF” contain the strong ribosome binding site (RBS) derived from phage T7 major capsid protein. The construct was verified by sequencing.

#### pGFPMutS

*gfpmut2* gene was PCR amplified using primers GFPmutSF and phosphorylated GFPmutSR. *ThemutS* gene was amplified using phosphorylated MutSF and EcoMutSR. These two fragments were ligated and resultant gene with in-frame GFP-MutS was re-amplified using primers GFPMutSF & EcoMutSR and cloned at Nhe-I and Sal-I sites of pBAD18kan. There is pro-gly-gly-serine linker between GFPmut2 and MutS for assistance in folding. The construct was verified by sequencing.Through complentation assays in *ΔmutS*, GFP-MutS functionality was confirmed.

### Colony size and generation number estimation

Before the start of mutation-accumulation experiments, a line of each genotype was grown to exponential stage in LB with 40μg/ml kanamycin and dilutions were spread on LB agar with kanamycin (40μg/ml) and incubated for 24 hours. Agar plugs containing colonies of various sizes (3mm to 5mm) for each genotype was suspended in 1ml of 1X M9 salt solution. Dilutions of this suspension were spread on LB agar with kanamycin and colonies were scored after 16 hours. Assuming that each colony arose from a single cell, the number of cells in a colony provides a measure to estimate the estimate the number of generations cell has divided to produce a colony of given size. In our study, across all genotype, on an average a colony sized 3.94mm corresponded to around 28.14 generations.

### Mutation accumulation experiment

As detailed earlier^24^, the mutation accumulation experiment was conducted. After transformation of appropriate plasmids in the parent 102BW *E.coli* strain, clonal cell lines of following genotypes were established: 9 control lines, 19 MutS overexpressing lines, 9 MutL overexpressing lines and 9 MutS-MutL expressing lines. Each line was passaged through single individual bottlenecks for 50 days on LB agar with kanamycin (40μg/ml). During the course of 50 days, at every 10^th^ day colony size was measured through a ruler. Additionally, every 10^th^day, the passaged colony was replica plated on LB agar with kanamycin (40μg/ml) and the next day, the cell were scrapped and suspended in 15% LB glycerol to create frozen stocks. At the end of experiment, the average size of colony for each genotype enabled the estimation of total number of generations elapsed for each genotype. At the end of 50 days, a colony from each line was patched on to a LB agar with kanamycin (40μg/ml) and post 24 hours, cells were scrapped and collected. From these cells, using Masterpure complete DNA and RNA purification kit (Lucigen), genomic DNA for whole genome sequencing was isolated. The parent strain 102BW which did not participate in the mutation accumulation experiment was also sequenced to weed out any mutations fixed prior to this experiment.

### Whole genome sequencing of mutation-accumulation lines

Truseq PCR free libraries were paired end (2 × 150bps) sequenced on NovaSeq-6000 platform using a S1 flow cell. For all samples, >1 million paired end reads were aligned against a circular reference genome of *E.coli* K12 strain BW25113 (GenBank Accession : CP009273.1). The average coverage for the entire experiment was around 60X. Using CLC genomics workbench 20.0.3, reads were initially mapped locally to reference using default settings and then subjected to local realignment process. SNP Variants which fulfilled either of the following conditions were called: (A) Frequency of alternate allele≥ 50%, Average Quality ≥ 30 and Minimum Alternate allele count ≥ 4. (B) Frequency of alternate allele≥ 10%, Average Quality ≥ 30, Minimum Alternate allele count ≥ 4 and Forward/reverse read ratio > 0. Small indels (≤ 4bps) were called only when frequency of alternate allele ≥ 50%, Average Quality ≥ 30 and Minimum Alternate allele count ≥ 10. Any variants present in repeat regions were omitted from the study. Large deletions were called using the “Advanced structural variant detection” tool at default settings. In MutS overexpression, all large deletions were confirmed using conventional sequencing. In case of MutS overexpression line no. 10, even though the “Advanced structural variant detection” tool did not show any deletions, while validating some falsely called MNVs in the repetitive regions, we confirmed a deletion at the repetitive element through conventional sequencing. The annotation of mutations and deletions was done manually using existing annotation databases.

### Inverting *lacZYA* orientation at its native locus

We inverted the entire *lacZYA* operon utilizing the FRT recombination system. Using *lacI::tet* as marker we brought the E461G-*lacZ*-allele in MG1655 background and named the strain as 102MG. We transduced *ΔlacI::Kan* from the keio mutant JW0336 into the 102MG and used pCP20 to flip out the kanamycin resistance cassette which left a scar sequence with a FRT site. To this strain, we transduced *ΔcynX::kan* from keio mutant JW0332. As a result of this there were three FRT sites in the genome, two similarly oriented FRT sites at*cynX::kan* and one opposite oriented FRT site at *lacI*::FRT. We again used pCP20 and screened for clones in which Flp recombinase had not only have removed antibiotic cassette but also caused the invertion of *lacZYA* operon. Through PCR, we selected two clones with invert orientation of *lacZYA* operon. As control, a clone with a native orientation of *lacZYA* operon with removed antibiotic resistance cassette was selected. Using PCR, it was confirmed that there was no duplication in the inverted region and junction were sequenced to confirm the inversion of *lacZYA* operon. The same was done for the clone with *lacZYA* operon in native orientation (Supplementary Fig.7 A-C).

### Generation of *E.coli* knockouts

We used linear dsDNA based recombineering in DY378 as describe^65^ to generate the required knockouts. For every knockout made in this study, two primers (each with around 30-40 bp homology to target gene and 22bp priming site) were used to amplify an antibiotic resistance cassette. The primer sequences can be found in the attached primer list.The template for this PCR was either pKD3 or an ampicillin resistance cassette containing the Amp promoter and β-lactamase gene (derived from pUC vector) flanked by the priming sites found in pKD3. The PCR product was gel purified, mixed with recombineering proficient electrocompetent DY378 cells and subsequently electrophorated. Immediately after electrophoration, cell were recovered in prewarmed 1ml SOC media (32°C) and incubated with shaking at 32°C for an hour. Afterwards, cells were pelleted, plated on appropriate antibiotic containing LB plates and incubated at 32°C. Genomic DNA from selected clones was isolated and PCR was conducted to confirm gene replacement. Of the two primers utilized for confirmation, one always annealed to the priming site in resistance cassette and the other upstream of the targeted region. P1 lysates were made on selected clones and the same was used to bring the mutation to 102BW strain. In case of Δ*mutH* and Δ*mutL* knockouts, junction regions where recombination of antibiotic

### ssDNA recombineering

*E.coli* Strain DY378 cells harboring either the control or pMutS or pF36AMutS plasmid were made recombineering proficient and electrocompetent as per the methods previously described^33^. Around 200ng of ssDNA oligo probe was mixed with 25ul of competent cells and pipetted into 0.1cm Biorad gene pulser cuvettes. Electrophoration was done using Biorad’s Genepulser dialled at 1.8KV, 25μF with pulse controller at 200ohms. Post electrophoration, 1ml of prewarmed (at 32°C) LB was immediately added and mixed gently by inverting the cuvette. Following this, the cells were diluted in LB with 40μg/ml kanamycin to a final volume of 10ml and were shaken at 32°C and 200rpm for 4.5 hours and subsequently aliquots of 1ml and 0.1ml were taken and spread on LB rifampicin plates and incubated at 32°C. Viable cell count was also measured by plating on LB agar. Plates were scored post 24 hrs and mutation frequency was determined.

### Microscopy

Most of the imaging was done on live cells. For Syto9 staining, cells at desired optical density were suspended in 0.85% saline. Then the cells were stained with Syto9 stain at a final concentration of 1μM and kept at 37°C water-bath for 10-15 minutes. For staining DNA with DAPI, the cells were fixed as per the protocol mentioned in earlier study^66^. To fix the cells, culture once grown till required, were mixed with equal volume of 5.2% paraformaldehyde, 0.012% glutaraldehyde, and 80 mM sodium phosphate (pH 7.5) and kept on ice for 5 to 10 minutes. The fixed cells were stained with 0.25μg/ml of DAPI in sodium phosphate buffer (pH 7.5) and kept at 37°C water-bath for 10-15 minutes. For microscopy, on a glass slide, cell suspensions were trapped in between cover slip and agarose pads. DIC and fluorescence images were taken with a 100X objective lens using Axioimager microscope (Carl Zeiss). Images were analyzed using Axiovision LE software.

### Immunoprecipitation

Cells grown in 5ml volume to desired optical density were suspended in 500μl cell lysis buffer (150mM NaCl, 0.5mM EDTA, 0.5% NP40, 1mM PMSF, 0.5mg/ml Lysozyme, cOmplete protease inhibitor[Roche], 10mM Tris pH 7.5).The suspension is kept on ice for 15-30 minutes on ice and then sonicated. The cell lysates were clarified by centrifuging at 15,000 rpm at 4°C for 30 minutes. The clarified lysated were mixed with GST tagged AntiGFP nanobody immobilized on glutathione agarose beads^67^. After 4 hours incubation in cold, the beads were collected and washed twice with PBS (phosphate buffer saline). The beads were mixed with 1X SDS-PAGE laemmli buffer, boiled and electrophoresed on 10% SDS-PAGE gel which was followed by western blotting. Membrane was probed using Anti-GFP and Anti-FtsZ antibody at dilution as described above in this section.

### Statiscal analysis

Chi-Square analysis were conducted using standard methods^68^. Using Kolmogorov-Smirnov Test of Normality, we confirmed that the mutations for each group in Mutation accumulation experiment followed a Normal Distribution. Two tailed t-tests were used for reporting results of mutation frequencies, β-galactosidase assaysand cell length analysis.

## Acknowledgements

We express deep gratitude to Dr.Manjula Reddy for her kind gift of strains, plasmids and scientific discussions. VRI thanks UGC for JRF & SRF fellowship for his research. SCM thanks CSIR JRF & SRF fellowship for her research. SK thanks Department of Biotechnology, India for partly funding this research.

## Author contributions

VRI conceived the study. VRI and SK designed the study. VRI and SCM executed the study. VRI, SCM and SK wrote the manuscript.

## Materials and correspondence

Correspondence and material requests can be addressed to the corresponding authors.

## References

1 Fijalkowska, I. J., Schaaper, R. M. & Jonczyk, P. DNA replication fidelity in Escherichia coli: a multi-DNA polymerase affair. FEMS microbiology reviews 36, 1105–1121, doi:10.1111/j.1574-6976.2012.00338.x (2012).

2 Modrich, P. DNA MISMATCH CORRECTION. Annual Review of Biochemistry 56, 435–466, doi:10.1146/annurev.bi.56.070187.002251 (1987).

3 Grilley, M., Welsh, K. M., Su, S. S. & Modrich, P. Isolation and characterization of the Escherichia coli mutL gene product. The Journal of biological chemistry 264, 1000–1004 (1989).

4 Au, K. G., Welsh, K. & Modrich, P. Initiation of methyl-directed mismatch repair. The Journal of biological chemistry 267, 12142–12148 (1992).

5 Pukkila, P. J., Peterson, J., Herman, G., Modrich, P. & Meselson, M. Effects of high levels of DNA adenine methylation on methyl-directed mismatch repair in Escherichia coli. Genetics 104, 571–582 (1983).

6 Marinus, M. G. DNA Mismatch Repair. EcoSal Plus 5, 10.1128/ecosalplus.1127.1122.1125, doi:10.1128/ecosalplus.7.2.5 (2012).

7 Bi, E. F. & Lutkenhaus, J. FtsZ ring structure associated with division in Escherichia coli. Nature 354, 161–164, doi:10.1038/354161a0 (1991).

8 de Boer, P., Crossley, R. & Rothfield, L. The essential bacterial cell-division protein FtsZ is a GTPase. Nature 359, 254–256, doi:10.1038/359254a0 (1992).

9 Margolin, W. FtsZ and the division of prokaryotic cells and organelles. Nat Rev Mol Cell Biol 6, 862–871, doi:10.1038/nrm1745 (2005).

10 Mukherjee, A., Cao, C. & Lutkenhaus, J. Inhibition of FtsZ polymerization by SulA, an inhibitor of septation in Escherichia coli. Proceedings of the National Academy of Sciences of the United States of America 95, 2885–2890, doi:10.1073/pnas.95.6.2885 (1998).

11 Huisman, O. & D’Ari, R. An inducible DNA replication-cell division coupling mechanism in E. coli. Nature 290, 797–799, doi:10.1038/290797a0 (1981).

12 Bi, E. & Lutkenhaus, J. Cell division inhibitors SulA and MinCD prevent formation of the FtsZ ring. Journal of bacteriology 175, 1118–1125, doi:10.1128/jb.175.4.1118-1125.1993 (1993).

13 Darmon, E., Eykelenboom, J. K., Lopez-Vernaza, M. A., White, M. A. & Leach, D. R. F. Repair on the Go: E. coli Maintains a High Proliferation Rate while Repairing a Chronic DNA Double-Strand Break. PLOS ONE 9, e110784, doi:10.1371/journal.pone.0110784 (2014).

14 White, M. A., Darmon, E., Lopez-Vernaza, M. A. & Leach, D. R. F. DNA double strand break repair in Escherichia coli perturbs cell division and chromosome dynamics. PLOS Genetics 16, e1008473, doi:10.1371/journal.pgen.1008473 (2020).

15 Gutierrez, A. et al. β-lactam antibiotics promote bacterial mutagenesis via an RpoS-mediated reduction in replication fidelity. Nature Communications 4, 1610, doi:10.1038/ncomms2607 (2013).

16 Tsui, H. C., Feng, G. & Winkler, M. E. Negative regulation of mutS and mutH repair gene expression by the Hfq and RpoS global regulators of Escherichia coli K-12. Journal of bacteriology 179, 7476, doi:10.1128/jb.179.23.7476-7487.1997 (1997).

17 Chen, J. & Gottesman, S. Hfq links translation repression to stress-induced mutagenesis in E. coli. Genes Dev 31, 1382–1395, doi:10.1101/gad.302547.117 (2017).

18 Nghiem, Y., Cabrera, M., Cupples, C. G. & Miller, J. H. The mutY gene: a mutator locus in Escherichia coli that generates G.C----T.A transversions. Proceedings of the National Academy of Sciences of the United States of America 85, 2709–2713, doi:10.1073/pnas.85.8.2709 (1988).

19 Cupples, C. G. & Miller, J. H. A set of lacZ mutations in Escherichia coli that allow rapid detection of each of the six base substitutions. Proceedings of the National Academy of Sciences of the United States of America 86, 5345–5349, doi:10.1073/pnas.86.14.5345 (1989).

20 Luria, S. E. & Delbrück, M. Mutations of Bacteria from Virus Sensitivity to Virus Resistance. Genetics 28, 491–511 (1943).

21 Garibyan, L. et al. Use of the rpoB gene to determine the specificity of base substitution mutations on the Escherichia coli chromosome. DNA repair 2, 593–608, doi:10.1016/s1568-7864(03)00024-7 (2003).

22 Miller, J. H. et al. Escherichia coli strains (ndk) lacking nucleoside diphosphate kinase are powerful mutators for base substitutions and frameshifts in mismatch-repair-deficient strains. Genetics 162, 5–13 (2002).

23 Schaaper, R. M. & Dunn, R. L. Spectra of spontaneous mutations in Escherichia coli strains defective in mismatch correction: the nature of in vivo DNA replication errors. Proceedings of the National Academy of Sciences of the United States of America 84, 6220–6224, doi:10.1073/pnas.84.17.6220 (1987).

24 Lee, H., Popodi, E., Tang, H. & Foster, P. L. Rate and molecular spectrum of spontaneous mutations in the bacterium Escherichia coli as determined by whole-genome sequencing. Proceedings of the National Academy of Sciences of the United States of America 109, E2774–2783, doi:10.1073/pnas.1210309109 (2012).

25 Foster, P. L., Lee, H., Popodi, E., Townes, J. P. & Tang, H. Determinants of spontaneous mutation in the bacterium Escherichia coli as revealed by whole-genome sequencing. Proceedings of the National Academy of Sciences of the United States of America 112, E5990–5999, doi:10.1073/pnas.1512136112 (2015).

26 Fijalkowska, I. J., Jonczyk, P., Tkaczyk, M. M., Bialoskorska, M. & Schaaper, R. M. Unequal fidelity of leading strand and lagging strand DNA replication on the *Escherichia coli* chromosome. Proceedings of the National Academy of Sciences 95, 10020, doi:10.1073/pnas.95.17.10020 (1998).

27 Maslowska, K. H., Makiela-Dzbenska, K., Mo, J. Y., Fijalkowska, I. J. & Schaaper, R. M. High-accuracy lagging-strand DNA replication mediated by DNA polymerase dissociation. Proceedings of the National Academy of Sciences of the United States of America l15, 42124217, doi:10.1073/pnas.1720353115 (2018).

28 Yamamoto, A., Schofield, M. J., Biswas, I. & Hsieh, P. Requirement for Phe36 for DNA binding and mismatch repair by Escherichia coli MutS protein. Nucleic acids research 28, 3564–3569, doi:10.1093/nar/28.18.3564 (2000).

29 López de Saro, F. J., Marinus, M. G., Modrich, P. & O’Donnell, M. The beta sliding clamp binds to multiple sites within MutL and MutS. The Journal of biological chemistry 281, 14340‐14349, doi:10.1074/jbc.M601264200 (2006).

30 Pluciennik, A., Burdett, V., Lukianova, O., O’Donnell, M. & Modrich, P. Involvement of the beta clamp in methyl-directed mismatch repair in vitro. The Journal of biological chemistry 284, 32782–32791, doi:10.1074/jbc.M109.054528 (2009).

31 Lu, A. L., Clark, S. & Modrich, P. Methyl-directed repair of DNA base-pair mismatches in vitro. Proceedings of the National Academy of Sciences of the United States of America 80, 4639–4643, doi:10.1073/pnas.80.15.4639 (1983).

32 Brown, J., Brown, T. & Fox, K. R. Affinity of mismatch-binding protein MutS for heteroduplexes containing different mismatches. The Biochemical journal 354, 627–633, doi:10.1042/0264-6021:3540627 (2001).

33 Ellis, H. M., Yu, D., DiTizio, T. & Court, D. L. High efficiency mutagenesis, repair, and engineering of chromosomal DNA using single-stranded oligonucleotides. Proceedings of the National Academy of Sciences of the United States of America 98, 6742–6746, doi:10.1073/pnas.121164898 (2001).

34 Iyer, R. R., Pluciennik, A., Burdett, V. & Modrich, P. L. DNA mismatch repair: functions and mechanisms. Chemical reviews 106, 302–323, doi:10.1021/cr0404794 (2006).

35 Bjedov, I. et al. Stress-induced mutagenesis in bacteria. Science (New York, N.Y.) 300, 1404–1409, doi:10.1126/science.1082240 (2003).

36 Cairns, J. & Foster, P. L. Adaptive reversion of a frameshift mutation in Escherichia coli. Genetics 128, 695–701 (1991).

37 Foster, P. L. Adaptive mutation in Escherichia coli. Cold Spring Harb Symp Quant Biol 65, 21–29, doi:10.1101/sqb.2000.65.21 (2000).

38 Robles, S. J. & Adami, G. R. Agents that cause DNA double strand breaks lead to p16INK4a enrichment and the premature senescence of normal fibroblasts. Oncogene 16, 1113–1123, doi:10.1038/sj.onc.1201862 (1998).

39 Povirk, L. F., Hogan, M. & Dattagupta, N. Binding of bleomycin to DNA: intercalation of the bithiazole rings. Biochemistry 18, 96–101, doi:10.1021/bi00568a015 (1979).

40 Pennington, J. M. & Rosenberg, S. M. Spontaneous DNA breakage in single living Escherichia coli cells. Nature genetics 39, 797–802, doi:10.1038/ng2051 (2007).

41 Huisman, O., D’Ari, R. & Gottesman, S. Cell-division control in Escherichia coli: specific induction of the SOS function SfiA protein is sufficient to block septation. Proceedings of the National Academy of Sciences of the United States of America 81, 4490–4494, doi:10.1073/pnas.81.14.4490 (1984).

42 Hill, S. A. & Little, J. W. Allele replacement in Escherichia coli by use of a selectable marker for resistance to spectinomycin: replacement of the lexA gene. Journal of bacteriology 170, 5913–5915, doi:10.1128/jb.170.12.5913-5915.1988 (1988).

43 SaiSree, L., Reddy, M. & Gowrishankar, J. lon incompatibility associated with mutations causing SOS induction: null uvrD alleles induce an SOS response in Escherichia coli. Journal of bacteriology 182, 3151–3157, doi:10.1128/jb.182.11.3151-3157.2000 (2000).

44 Bernhardt, T. G. & de Boer, P. A. SlmA, a nucleoid-associated, FtsZ binding protein required for blocking septal ring assembly over Chromosomes in E. coli. Mol Cell 18, 555–564, doi:10.1016/j.molcel.2005.04.012 (2005).

45 Zhao, J. & Winkler, M. E. Reduction of GC → TA Transversion Mutation by Overexpression of MutS in *Escherichia coli* K-12. Journal of bacteriology 182, 5025–5028, doi:10.1128/jb.182.17.5025-5028.2000 (2000).

46 Qiu, R. et al. MutL traps MutS at a DNA mismatch. Proceedings of the National Academy of Sciences 112, 10914, doi:10.1073/pnas.1505655112 (2015).

47 Hall, M. C. & Matson, S. W. The Escherichia coli MutL protein physically interacts with MutH and stimulates the MutH-associated endonuclease activity. The Journal of biological chemistry 274, 1306–1312, doi:10.1074/jbc.274.3.1306 (1999).

48 Sano, E., Maisnier-Patin, S., Aboubechara, J. P., Quiñones-Soto, S. & Roth, J. R. Plasmid copy number underlies adaptive mutability in bacteria. Genetics 198, 919–933, doi:10.1534/genetics.114.170068 (2014).

49 Bernhardt, T. G. & de Boer, P. A. Screening for synthetic lethal mutants in Escherichia coli and identification of EnvC (YibP) as a periplasmic septal ring factor with murein hydrolase activity. Molecular microbiology 52, 1255–1269, doi:10.1111/j.1365-2958.2004.04063.x (2004).

50 Vinella, D., Joseleau-Petit, D., Thévenet, D., Bouloc, P. & D’Ari, R. Penicillin-binding protein 2 inactivation in Escherichia coli results in cell division inhibition, which is relieved by FtsZ overexpression. Journal of bacteriology 175, 6704–6710, doi:10.1128/jb.175.20.6704-6710.1993 (1993).

51 Chakraborty, U., Dinh, T. A. & Alani, E. Genomic Instability Promoted by Overexpression of Mismatch Repair Factors in Yeast: A Model for Understanding Cancer Progression. 209, 439–456, doi:10.1534/genetics.118.300923 (2018).

52 Yu, J., Liu, Y., Yin, H. & Chang, Z. Regrowth-delay body as a bacterial subcellular structure marking multidrug-tolerant persisters. Cell Discovery 5, 8, doi:10.1038/s41421-019-0080-3 (2019).

53 Kauffmann, A. et al. High expression of DNA repair pathways is associated with metastasis in melanoma patients. Oncogene 27, 565–573, doi:10.1038/sj.onc.1210700 (2008).

54 Wilczak, W. et al. Up-regulation of mismatch repair genes MSH6, PMS2 and MLH1 parallels development of genetic instability and is linked to tumor aggressiveness and early PSA recurrence in prostate cancer. Carcinogenesis 38, 19–27, doi:10.1093/carcin/bgw116 (2017).

55 Wagner, V. P. et al. Overexpression of MutSα Complex Proteins Predicts Poor Prognosis in Oral Squamous Cell Carcinoma. Medicine 95, e3725, doi:10.1097/md.0000000000003725 (2016).

56 Albero-González, R. & Hernández-Llodrà, S. Immunohistochemical expression of mismatch repair proteins (MSH2, MSH6, MLH1, and PMS2) in prostate cancer: correlation with grade groups (WHO 2016) and ERG and PTEN status. 475, 223–231, doi:10.1007/s00428-019-02591-z (2019).

57 Alexandrov, L. B. et al. Signatures of mutational processes in human cancer. Nature 5OO, 415–421, doi:10.1038/nature12477 (2013).

58 Dufner, P., Marra, G., Räschle, M. & Jiricny, J. Mismatch recognition and DNA-dependent stimulation of the ATPase activity of hMutSalpha is abolished by a single mutation in the hMSH6 subunit. The Journal of biological chemistry 275, 36550–36555, doi:10.1074/jbc.M005987200 (2000).

59 Bowers, J., Sokolsky, T., Quach, T. & Alani, E. A mutation in the MSH6 subunit of the Saccharomyces cerevisiae MSH2-MSH6 complex disrupts mismatch recognition. The Journal of biological chemistry 274, 16115–16125, doi:10.1074/jbc.274.23.16115 (1999).

60 Khlebnikov, A., Datsenko, K. A., Skaug, T., Wanner, B. L. & Keasling, J. D. Homogeneous expression of the P(BAD) promoter in Escherichia coli by constitutive expression of the low-affinity high-capacity AraE transporter. Microbiology (Reading, England) 147, 3241–3247, doi:10.1099/00221287-147-12-3241 (2001).

61 Miller, J. H. A short course in bacterial genetics: a laboratory manual and handbook for Escherichia coli and related bacteria. (New York (N.Y.): Cold Spring Harbor laboratory press, 1992).

62 Gillet-Markowska, A., Louvel, G. & Fischer, G. *bz-rates*: A Web Tool to Estimate Mutation Rates from Fluctuation Analysis. G3: Genes|Genomes|Genetics 5, 2323–2327, doi:10.1534/g3.115.019836 (2015).

63 Kumar, S., Stecher, G., Li, M., Knyaz, C. & Tamura, K. MEGA X: Molecular Evolutionary Genetics Analysis across Computing Platforms. Molecular biology and evolution 35, 1547–1549, doi:10.1093/molbev/msy096 (2018).

64 Guzman, L. M., Belin, D., Carson, M. J. & Beckwith, J. Tight regulation, modulation, and high-level expression by vectors containing the arabinose PBAD promoter. Journal of bacteriology 177, 4121–4130, doi:10.1128/jb.177.14.4121-4130.1995 (1995).

65 Yu, D. et al. An efficient recombination system for chromosome engineering in *Escherichia coli*. Proceedings of the National Academy of Sciences 97, 5978, doi:10.1073/pnas.100127597 (2000).

66 Pogliano, J., Pogliano, K., Weiss, D. S., Losick, R. & Beckwith, J. Inactivation of FtsI inhibits constriction of the FtsZ cytokinetic ring and delays the assembly of FtsZ rings at potential division sites. Proceedings of the National Academy of Sciences 94, 559–564, doi:10.1073/pnas.94.2.559 (1997).

67 Katoh, Y., Nozaki, S., Hartanto, D., Miyano, R. & Nakayama, K. Architectures of multisubunit complexes revealed by a visible immunoprecipitation assay using fluorescent fusion proteins. Journal of Cell Science 128, 2351, doi:10.1242/jcs.168740 (2015).

68 Zar, J. H. Biostatistical Analysis (5th Edition). (Prentice-Hall, Inc., 2007).

