## Supplementary Figure 1 to 7 for "Overexpression of MutS impairs DNA mismatch repair and causes cell division defect in *E.coli*"

Supplementary fig 1

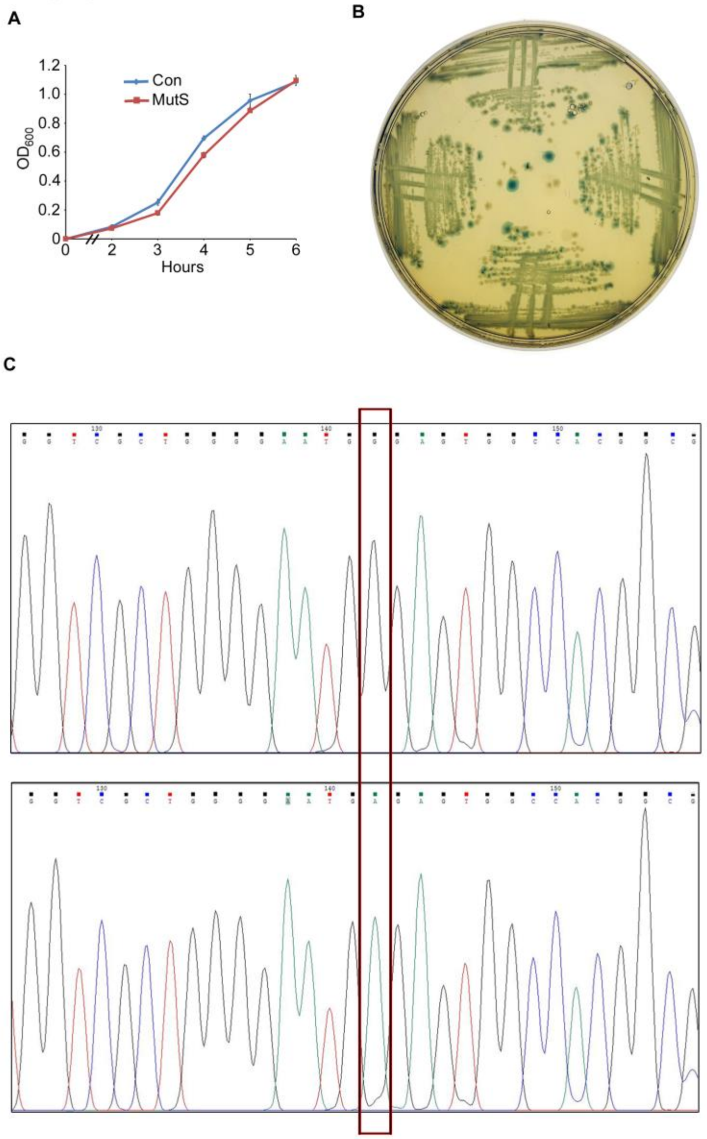


**Supplementary figure 1: (A)** Growth curves of 102BW E.coli cells carrying either vector control (Con) or pMutS (MutS). Overnight cultures were diluted 1: 1000 in fresh media and optical density at 600nm was measured at indicated time points and plotted. Each data points represent average of three independent experiments and error bar represent ±SD. **(B)** Representative image of streaked blue papillae. The blue papillae observed 48 hrs after streaking of pMutS carrying 102BW E.coli cells on LB-Kan X-gal Lactose plate, were picked and streaked on to a fresh LB-Kan X-gal Lactose plate. After overnight incubation, resultant blue streaks were streaked on to a fresh LB-Kan X-gal Lactose plates to enrich the blue colonies. **(C)** Chromatogram showing the sequence of a region of E461G lacZ allele before (top panel) and after undergoing GC to AT mutation (lower panel). The mutated sequence is derived from the sequencing of the *lacZ* region from the blue colonies. The critical 1385$^{th}$ nucleotide of E461G *lacZ* allele which had undergone a GC to AT mutation and restores lac$^{+}$ phenotype is marked with the help of a red rectangle.

**Supplementary fig 2**

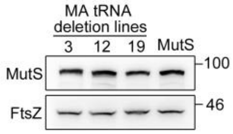


**Supplementary figure 2:** Western blot showing MutS protein levels in exponential phase cells of MutS-overexpressing mutation accumulation (MA) lines, exhibiting deletion of intergenic sequences in tRNA locus after ≈1250 generations. Line No.s 3, 12 and 19 poses a deletion of intergenic sequences between *glnX* and *glnV*, *tyrV* and *tyrX* and *alaX* and *alaW* tRNA genes respectively. “Isogenic strain” represents exponential phased cells of 102 BW cells carrying pMutS (MutS) which had not participated in the MA experiment.

**Supplementary fig 3**

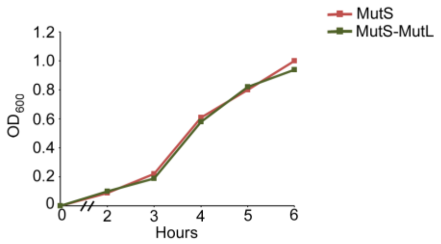


**Supplementary figure 3:** Growth curves of 102BW E.coli cells carrying either pMutS (MutS) or pMutS-MutL (MutS-MutL). Overnight cultures were diluted 1: 1000 in fresh media and optical density at 600nm was measured at indicated time points and plotted.

**Supplementary fig 4**

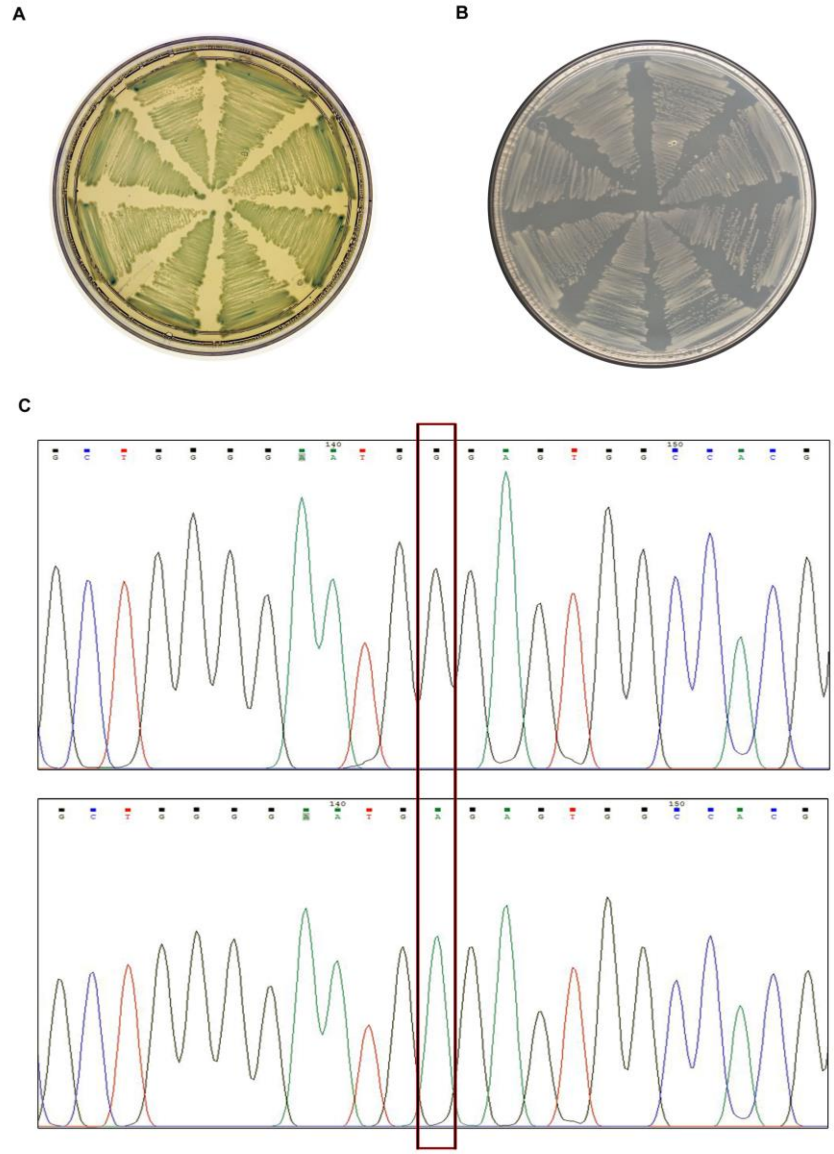


**Supplementary figure 4:** Representative image of streaked Lac$^{+}$ revertants on **(A)** LB-Kan X-gal Lactose **(B)** Minimal lactose plates. **(C)** Chromatogram showing the sequence of a region of E461G lacZ allele before (top panel) and after undergoing GC to AT mutation (lower panel). The mutated sequence is derived from the sequencing of the *lacZ* region from the Lac$^{+}$ revertants. The critical 1385$^{th}$ nucleotide of E461G *lacZ* allele which had undergone a GC to AT mutation and restores lac$^{+}$ phenotype is marked with the help of a red rectangle.

Supplementary fig 5

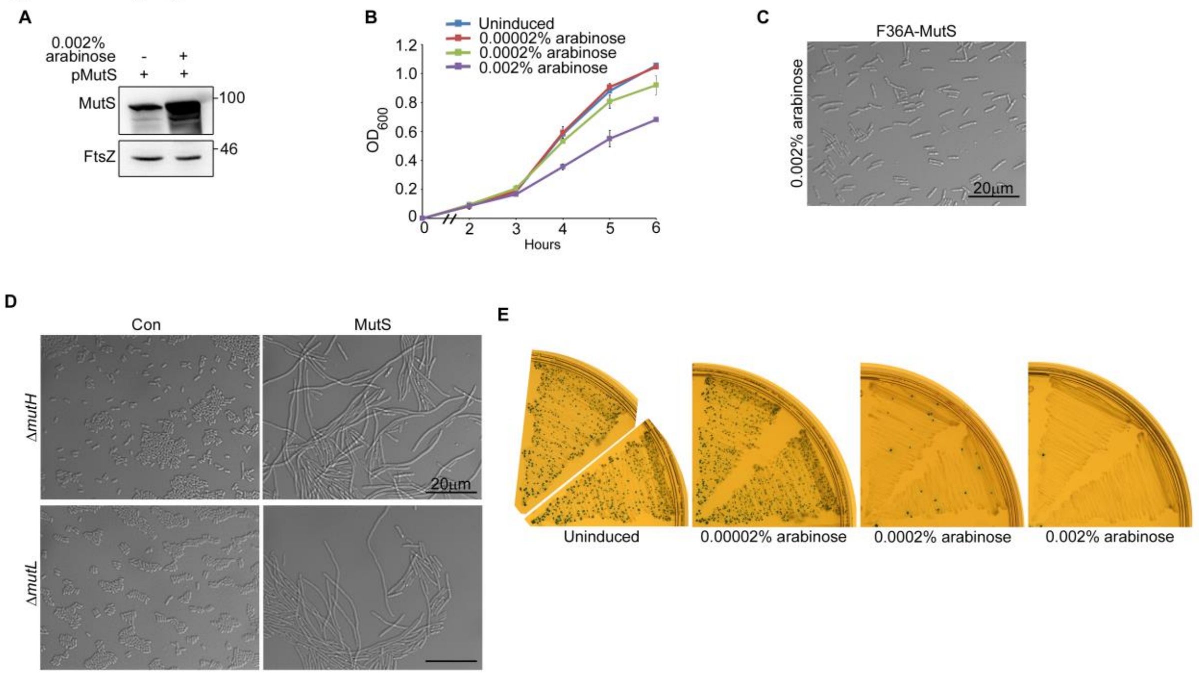


**Supplementary figure 5: (A)**Western blot showing MutS expression levels in exponential phase 102 BW *E.coli* cells carrying pMutS when grown under uninduced and 0.002% arabinose induced conditions **(B)** Growth curves of 102BW *E.coli* cells carrying pMutS (MutS) grown without and with the indicated concentrations of arabinose. Overnight culture of *E.coli* cells was diluted 1:1000 in fresh media, shaken till $OD_{600}\approx 0.1$ and induced with arabinose (concentration as indicated) and optical density at 600nm was measured at indicated time points and plotted. Each data points represent average of three independent experiments and error bar represent ±SD. **(C)** Representative DIC microscopy image of pF36AMutS carrying 102 BW cells. Overnight culture of pF36AMutS containing 102BW *E.coli* cells was diluted 1:1000 in fresh media, shaken till $OD_{600}\approx 0.1$, induced with 0.002% arabinose and grown further till $OD_{600}$= 0.6-0.7. **(D)** Representative DIC microscopy image of Δ*mutL* or Δ*mutH* 102BW *E.coli* cells carrying either vector control (Con) or pMutS (MutS).Overnight culture of cells was diluted 1:1000 in fresh media, shaken till $OD_{600}\approx 0.1$, induced with 0.002% arabinose and grown further till $OD_{600}$= 0.6-0.7. **(E)** Representative images of lac papillation assay in E461G *lacZ* allele. A single colony of 102BW cells carrying pMutS (MutS) was streaked on LB-Kan Xgal Lactose plates without and with indicated concentration of arabinose. Two replicates are shown for each condition. The number of blue papillae provides an estimate of the GC to AT mutation rate. Representative images are selected out of at least three independent experiments.

Supplementary fig 6

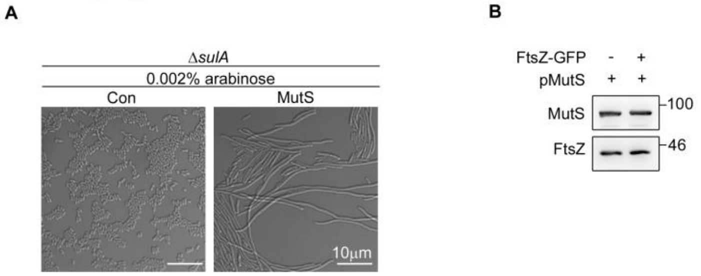


**Supplementary figure 6: (A)** Representative DIC microscopy image of Δ*sulA* 102BW *E.coli* cells carrying either vector control (Con) or pMutS (MutS). Overnight culture of cells was diluted 1:1000 in fresh media, shaken till $OD_{600}$≈0.1, induced with 0.002% arabinose and grown further till $OD_{600}$= 0.6-0.7. **(B)** Western blot showing MutS expression levels in exponential phased, pMutS carrying 102 BW or VRI7 *E.coli* cells. Representative images are selected out of at least three independent experiments.

### Supplementary fig 7

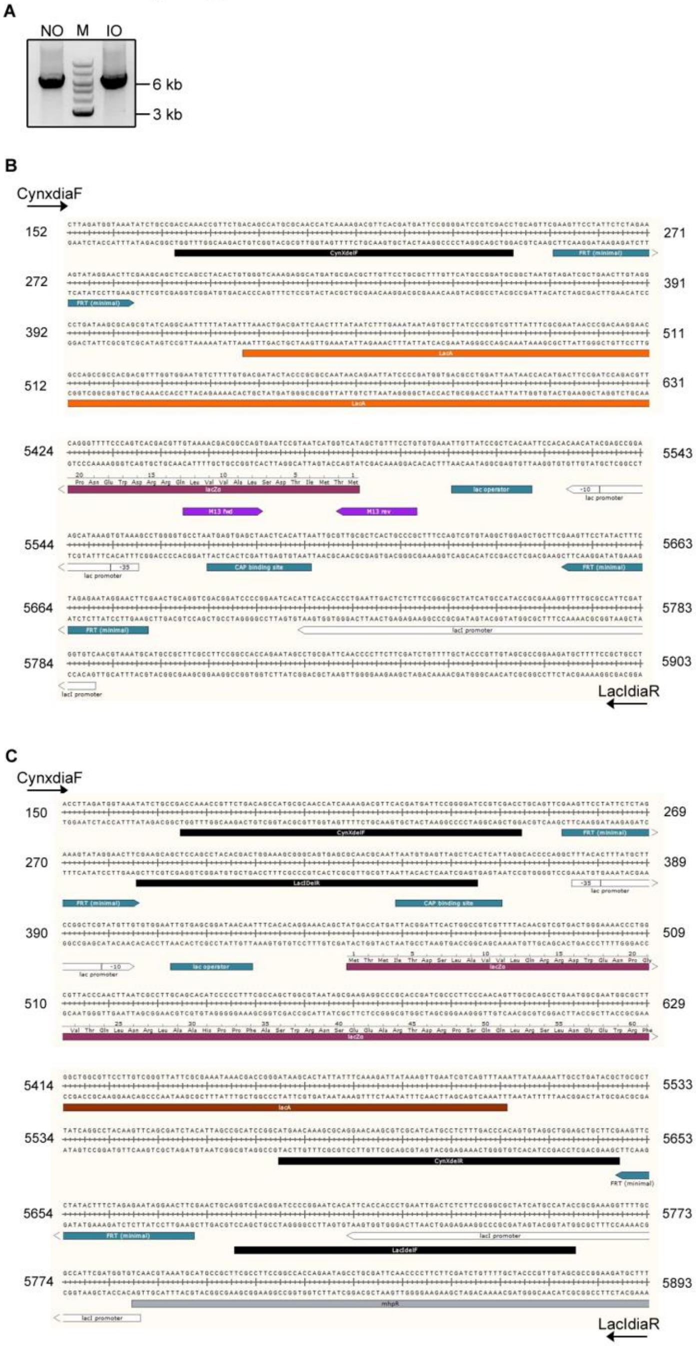


**Supplementary figure 7: (A)** Image depicts the 6.087 Kbp product of PCR amplification of entire *laczya* operon from MG1655 *E.coli* cells with E461G *lacZ* allele in either native orientation (NO) or invert orientation (IO). Using primers CynXdiaF and LacIdiaR, the entire region was amplified. (**B,C**) Structure and sequence of junction regions containing FRT sites in MG1655 *E.coli* cells with E461G *lacZ* allele in either **(B)** native orientation or **(C)** invert orientation. Primers CynXdiaF and LacIdiaR were utilized for sequencing the depicted regions.
