## Supplementary table 1 for "Overexpression of MutS impairs DNA mismatch repair and causes cell division defect in *E.coli*"

| **S.No.**  **Supplementary Table 1: Cumulative BPS spectrum in cluster-II region of *rpoB* gene conferring rifampicin resistance in 102BW *E.coli* cells overexpressing the indicated proteins** | **Nulceotide position in *rpoB* gene** | **Amino Acid change** | **Base pair substitution (BPS)** | **Control^a^** | **∆*mutS^a^* Control** | **MutS^*^ Overexpression** | **F36A MutS^*^ Overexpression** | **MutS-MutL^*^ Co-overexpression** |
| --- | --- | --- | --- | --- | --- | --- | --- | --- |
| 1 | 1522 | S508P | AT → GC | 0 | 0 | 0 | 0 | 0 |
| 2 | 1532 | L511P | AT → GC | 5 | 2 | 3 | 0 | 4 |
| 3 | 1534 | S512P | AT → GC | 2 | 0 | 8 | 0 | 0 |
| 4 | 1538 | Q513R | AT → GC | 0 | 1 | 4 | 1 | 2 |
| 5 | 1547 | D516G | AT → GC | 26 | 58 | 67 | 3 | 52 |
| 6 | 1552 | N518D | AT → GC | 0 | 0 | 1 | 0 | 0 |
| 7 | 1577 | H526R | AT → GC | 0 | 0 | 0 | 0 | 0 |
| 8 | 1598 | L533P | AT → GC | 4 | 0 | 2 | 4 | 4 |
| 9 | 1715 | I572T | AT → GC | 0 | 0 | 0 | 0 | 0 |
| 10 | 1520 | G507D | GC → AT | 0 | 0 | 0 | 0 | 0 |
| 11 | 1535 | S512F | GC → AT | 5 | 0 | 2 | 1 | 9 |
| 12 | 1546 | D516N | GC → AT | 2 | 24 | 27 | 1 | 5 |
| 13 | 1565 | S522F | GC → AT | 10 | 0 | 6 | 12 | 35 |
| 14 | 1576 | H526Y | GC → AT | 19 | 4 | 42 | 14 | 24 |
| 15 | 1585^ts^ | R529C | GC → AT | 3 | 0 | 2 | 0 | 2 |
| 16 | 1586 | R529H | GC → AT | 3 | 0 | 2 | 2 | 2 |
| 17 | 1592 | S531F | GC → AT | 17 | 1 | 13 | 13 | 13 |
| 18 | 1595 | A532V | GC → AT | 0 | 0 | 0 | 0 | 0 |
| 19 | 1600 | G534S | GC → AT | 0 | 0 | 0 | 0 | 0 |
| 20 | 1601 | G534D | GC → AT | 0 | 0 | 0 | 0 | 0 |
| 21 | 1691 | P564L | GC → AT | 10 | 1 | 15 | 13 | 7 |
| 22 | 1721 | S574F | GC → AT | 1 | 0 | 0 | 0 | 1 |
| 23 | 2060^ts^ | R687H | GC → AT | 0 | 0 | 0 | 0 | 0 |
| 24 | 1532 | L511Q | AT → TA | 0 | 0 | 0 | 0 | 0 |
| 25 | 1538 | Q513L | AT → TA | 25 | 0 | 7 | 13 | 12 |
| 26 | 1547 | D516V | AT → TA | 0 | 0 | 0 | 2 | 5 |
| 27 | 1568 | E523V | AT → TA | 0 | 0 | 0 | 0 | 0 |
| 28 | 1577 | H526L | AT → TA | 2 | 0 | 3 | 4 | 1 |
| 29 | 1598 | L533H | AT → TA | 0 | 0 | 0 | 0 | 0 |
| 30 | 1714 | I572F | AT → TA | 34 | 0 | 1 | 77 | 3 |
| 31 | 1525 | S509R | AT → CG | 4 | 0 | 0 | 0 | 0 |
| 32 | 1532 | L511R | AT → CG | 5 | 0 | 0 | 0 | 1 |
| 33 | 1534 | S512A | AT → CG | 0 | 0 | 0 | 0 | 0 |
| 34 | 1538 | Q513P | AT → CG | 7 | 0 | 1 | 1 | 6 |
| 35 | 1547 | D516A | AT → CG | 0 | 0 | 0 | 0 | 0 |
| 36 | 1577 | H526P | AT → CG | 3 | 0 | 0 | 0 | 0 |
| 37 | 1598 | L533R | AT → CG | 0 | 0 | 0 | 0 | 0 |
| 38 | 1687^ts^ | T563P | AT → CG | 0 | 0 | 0 | 0 | 0 |
| 39 | 1714 | I572L | AT → CG | 8 | 0 | 2 | 0 | 7 |
| 40 | 1715 | I572S | AT → CG | 21 | 0 | 3 | 0 | 1 |
| 41 | 1527 | S509R | GC → TA | 0 | 0 | 0 | 0 | 1 |
| 42 | 1535 | S512Y | GC → TA | 1 | 0 | 0 | 8 | 0 |
| **S.No.** | **Nulceotide position in *rpoB* gene** | **Amino Acid change** | **Base pair substitution (BPS)** | **Control^a^** | **∆*mutS^a^* Control** | **MutS^*^ Overexpression** | **F36A MutS^*^ Overexpression** | **MutS-MutL^*^ Co-overexpression** |
| 43 | 1537 | Q513K | GC → TA | 1 | 0 | 0 | 0 | 1 |
| 44 | 1546 | D516Y | GC → TA | 4 | 0 | 2 | 0 | 1 |
| 45 | 1565 | S522Y | GC → TA | 0 | 0 | 0 | 0 | 1 |
| 46 | 1576 | H526N | GC → TA | 5 | 1 | 17 | 9 | 0 |
| 47 | 1578 | H526Q | GC → TA | 1 | 0 | 0 | 0 | 0 |
| 48 | 1585^ts^ | R529S | GC → TA | 0 | 0 | 0 | 0 | 0 |
| 49 | 1586 | R529L | GC → TA | 0 | 0 | 0 | 0 | 0 |
| 50 | 1592 | S531Y | GC → TA | 0 | 0 | 0 | 0 | 14 |
| 51 | 1595 | A532E | GC → TA | 0 | 0 | 0 | 0 | 0 |
| 52 | 1600 | G534C | GC → TA | 0 | 0 | 0 | 0 | 0 |
| 53 | 1601 | G534V | GC → TA | 0 | 0 | 0 | 0 | 0 |
| 54 | 1708 | G570C | GC → TA | 0 | 0 | 0 | 0 | 0 |
| 55 | 1721 | S574Y | GC → TA | 0 | 0 | 0 | 0 | 0 |
| 57 | 1527 | S509R | GC → CG | 0 | 0 | 0 | 0 | 0 |
| 58 | 1574 | T525R | GC → CG | 0 | 0 | 0 | 0 | 0 |
| 59 | 1576 | H526D | GC → CG | 4 | 0 | 5 | 5 | 15 |
| 60 | 1578 | H526Q | GC → CG | 0 | 0 | 0 | 0 | 0 |
| 61 | 1601 | G534A | GC → CG | 0 | 0 | 0 | 0 | 0 |
| 62 | 1691 | P564R | GC → CG | 0 | 0 | 0 | 0 | 0 |
| 63^b,c^ | 1716 | I572M | GC → CG | 0 | 0 | 0 | 0 | 0 |
| **TOTAL** |  |  |  | **232** | **92** | **235** | **183** | **229** |
| Transition Mutations (Ts) | | |  | 107 | 91 | 194 | 64 | 160 |
| Transversion Mutations (Tv) | | |  | 125 | 1 | 41 | 119 | 69 |
| % of Transition Mutations | | |  | 46.12 | 98.91 | 82.55 | 34.97 | 69.87 |
| % of Transversion Mutations | | |  | 53.88 | 1.09 | 17.45 | 65.03 | 30.13 |
| Ts/Tv ratio | |  |  | 0.86 | 91.00 | 4.73 | 0.54 | 2.32 |
| Fold change in Ts/Tv ratio | | |  | 1.00 | 106.31 | 5.53 | 0.63 | 2.71 |

**^*^ pMutS, pF36AMutS and pMutS-MutL plasmids were used for overexpression of MutS, F36AMutS or MutS-MutL in 102BW E.coli cells**

**^a^ Empty vector control; pBAD18-Kan in either 102BW cells or VRI1 cells (∆*mutS* derivative of 102BW *E.coli* cells)**

**^b,c^There are total 71 sites which confer rifampicin resistance. In this study we have only analyzed 68 sites which lie in cluster-II region of *rpoB* gene**

**^b^ In our study, we also found a new mutation 1537 GC to CG in rpoB gene conferring rifampicin resistance**

**^ts^ Nucleotide position designated with the superscript “ts” confers temperature sensitive rifampicin resistance.**
