## Supplementary table 2 for "Overexpression of MutS impairs DNA mismatch repair and causes cell division defect in *E.coli*"

| **S.No.**  **Supplementary Table 2: Cumulative BPS spectrum in cluster-II region of *rpoB* gene conferring rifampicin**  **resistance in 102BW *E.coli* cells carrying pNRBSMutS (grown without or with indicated concentration of arabinose)** | **Nulceotide position in *rpoB* gene** | **Amino Acid change** | **Base pair substitution (BPS)** | **nRBS-MutS Uninduced** | **nRBS-MutS**  **induced with 0.0002% Arabinose** | **nRBS-MutS induced with 0.002% Arabinose** |
| --- | --- | --- | --- | --- | --- | --- |
| 1 | 1522 | S508P | AT → GC | 0 | 0 | 0 |
| 2 | 1532 | L511P | AT → GC | 0 | 1 | 3 |
| 3 | 1534 | S512P | AT → GC | 0 | 1 | 2 |
| 4 | 1538 | Q513R | AT → GC | 0 | 1 | 3 |
| 5 | 1547 | D516G | AT → GC | 16 | 10 | 36 |
| 6 | 1552 | N518D | AT → GC | 0 | 0 | 0 |
| 7 | 1577 | H526R | AT → GC | 1 | 0 | 0 |
| 8 | 1598 | L533P | AT → GC | 2 | 2 | 4 |
| 9 | 1715 | I572T | AT → GC | 0 | 0 | 0 |
| 10 | 1520 | G507D | GC → AT | 0 | 0 | 0 |
| 11 | 1535 | S512F | GC → AT | 1 | 1 | 3 |
| 12 | 1546 | D516N | GC → AT | 0 | 29 | 31 |
| 13 | 1565 | S522F | GC → AT | 6 | 11 | 2 |
| 14 | 1576 | H526Y | GC → AT | 25 | 10 | 7 |
| 15 | 1585^ts^ | R529C | GC → AT | 0 | 1 | 1 |
| 16 | 1586 | R529H | GC → AT | 1 | 0 | 1 |
| 17 | 1592 | S531F | GC → AT | 10 | 1 | 4 |
| 18 | 1595 | A532V | GC → AT | 0 | 0 | 0 |
| 19 | 1600 | G534S | GC → AT | 0 | 0 | 0 |
| 20 | 1601 | G534D | GC → AT | 0 | 0 | 0 |
| 21 | 1691 | P564L | GC → AT | 2 | 11 | 7 |
| 22 | 1721 | S574F | GC → AT | 1 | 0 | 0 |
| 23 | 2060^ts^ | R687H | GC → AT | 0 | 0 | 0 |
| 24 | 1532 | L511Q | AT → TA | 0 | 0 | 0 |
| 25 | 1538 | Q513L | AT → TA | 2 | 22 | 0 |
| 26 | 1547 | D516V | AT → TA | 12 | 1 | 1 |
| 27 | 1568 | E523V | AT → TA | 0 | 0 | 0 |
| 28 | 1577 | H526L | AT → TA | 0 | 0 | 1 |
| 29 | 1598 | L533H | AT → TA | 0 | 0 | 0 |
| 30 | 1714 | I572F | AT → TA | 0 | 1 | 4 |
| 31 | 1525 | S509R | AT → CG | 0 | 0 | 0 |
| 32 | 1532 | L511R | AT → CG | 0 | 0 | 0 |
| 33 | 1534 | S512A | AT → CG | 0 | 0 | 0 |
| 34 | 1538 | Q513P | AT → CG | 6 | 2 | 0 |
| 35 | 1547 | D516A | AT → CG | 0 | 0 | 0 |
| 36 | 1577 | H526P | AT → CG | 10 | 0 | 0 |
| 37 | 1598 | L533R | AT → CG | 0 | 0 | 0 |
| 38 | 1687^ts^ | T563P | AT → CG | 0 | 0 | 0 |
| 39 | 1714 | I572L | AT → CG | 11 | 1 | 1 |
| 40 | 1715 | I572S | AT → CG | 0 | 0 | 0 |
| 41 | 1527 | S509R | GC → TA | 0 | 0 | 0 |
| **S.No.** | **Nulceotide position in *rpoB* gene** | **Amino Acid change** | **Base pair substitution (BPS)** | **nRBS-MutS Uninduced** | **nRBS-MutS**  **induced with 0.0002% Arabinose** | **nRBS-MutS induced with 0.002% Arabinose** |
| 42 | 1535 | S512Y | GC → TA | 1 | 0 | 0 |
| 43 | 1537 | Q513K | GC → TA | 8 | 0 | 0 |
| 44 | 1546 | D516Y | GC → TA | 0 | 0 | 2 |
| 45 | 1565 | S522Y | GC → TA | 0 | 0 | 0 |
| 46 | 1576 | H526N | GC → TA | 5 | 11 | 3 |
| 47 | 1578 | H526Q | GC → TA | 0 | 0 | 0 |
| 48 | 1585^ts^ | R529S | GC → TA | 0 | 0 | 0 |
| 49 | 1586 | R529L | GC → TA | 0 | 0 | 0 |
| 50 | 1592 | S531Y | GC → TA | 0 | 0 | 0 |
| 51 | 1595 | A532E | GC → TA | 0 | 0 | 0 |
| 52 | 1600 | G534C | GC → TA | 0 | 0 | 0 |
| 53 | 1601 | G534V | GC → TA | 0 | 0 | 0 |
| 54 | 1708 | G570C | GC → TA | 0 | 0 | 0 |
| 55 | 1721 | S574Y | GC → TA | 0 | 0 | 0 |
| 57 | 1527 | S509R | GC → CG | 0 | 0 | 0 |
| 58 | 1574 | T525R | GC → CG | 0 | 0 | 0 |
| 59 | 1576 | H526D | GC → CG | 2 | 0 | 1 |
| 60 | 1578 | H526Q | GC → CG | 0 | 0 | 0 |
| 61 | 1601 | G534A | GC → CG | 0 | 0 | 0 |
| 62 | 1691 | P564R | GC → CG | 0 | 0 | 0 |
| 63^a^ | 1716 | I572M | GC → CG | 0 | 0 | 0 |
| **TOTAL** |  |  |  | 122 | 117 | 117 |
| Transition Mutations (Ts) | | |  | 65 | 79 | 104 |
| Transversion Mutations (Tv) | | |  | 57 | 38 | 13 |
| **% of Transition Mutations** | | |  | **53.28** | **67.52** | **88.89** |
| % of Transversion Mutations | | |  | 46.72 | 32.48 | 11.11 |
| **Ts/Tv ratio** | |  |  | **1.14** | **2.08** | **8.00** |
| **Fold change in Ts/Tv ratio** | | |  | **1.00** | **1.82** | **7.02** |

**^a^There are total 71 sites which confer rifampicin resistance. In this study we have only analyzed 68 sites which lie in cluster-II region of *rpoB* gene**

**^ts^ Nucleotide position designated with the superscript “ts” confers temperature sensitive rifampicin resistance.**
