## Supplementary table 3 for "Overexpression of MutS impairs DNA mismatch repair and causes cell division defect in *E.coli*"

| **Supplementary Table 3: Detailed outcome of Mutation Accumulation experiment in 102BW *E.coli* cells overexpressing the indicated proteins.** | **Control** | **Fraction** | **MutS overexpression** | **Fraction** | **MutL overexpresion** | **Fraction** | **MutS-MutL overxpression** | **Fraction** |
| --- | --- | --- | --- | --- | --- | --- | --- | --- |
| **SNPs** |  |  |  |  |  |  |  |  |
| Transitions | 9 | 0.5 | 18 | 0.64 | 12 | 0.55 | 4 | 0.26 |
| A:T to G:C | 3 | 0.17 | 6 | 0.21 | 4 | 0.18 | 2 | 0.13 |
| G:C to A:T | 6 | 0.33 | 12 | 0.43 | 8 | 0.36 | 2 | 0.13 |
| Transversions | 9 | 0.5 | 10 | 0.36 | 10 | 0.45 | 11 | 0.74 |
| A:T to C:G | 4 | 0.22 | 2 | 0.07 | 4 | 0.18 | 1 | 0.07 |
| A:T to T:A | 1 | 0.06 | 2 | 0.07 | 0 | 0 | 2 | 0.13 |
| G:C to T:A | 2 | 0.11 | 2 | 0.07 | 1 | 0.05 | 5 | 0.33 |
| G:C to C:G | 2 | 0.11 | 4 | 0.14 | 5 | 0.23 | 3 | 0.2 |
| AT sites | 8 | 0.44 | 10 | 0.36 | 8 | 0.36 | 5 | 0.33 |
| GC sites | 10 | 0.56 | 18 | 0.64 | 14 | 0.64 | 10 | 0.67 |
| Total | 18 |  | 28 |  | 22 |  | 15 |  |
| **Ts/Tv ratio** | **1** |  | **1.8** |  | **1.2** |  | **0.36** |  |
| **INDELS** | | | | | | | | |
| Small Indels (≤4 Base pairs) | | | | | | | | |
| Single nucleotide Indels | 0 | 0 | 5 | 0.36 | 2 | 0.5 | 0 | 0 |
| Multiple nucleotide Indels | 1 | 1 | 1 | 0.07 | 0 |  | 0 | 0 |
| Large Deletions | 0 | 0 | 8 | 0.57 | 2 | 0.5 | 0 | 0 |
| **Total** | **1** |  | **14** |  | **4** |  | **0** |  |
| **OUTCOME OF SNPs** | | | | | | | | |
| Positions |  |  |  |  |  |  |  |  |
| Coding | 13 | 0.72 | 24 | 0.86 | 19 | 0.86 | 13 | 0.87 |
| Non-coding | 5 | 0.28 | 4 | 0.14 | 3 | 0.14 | 1 | 0.07 |
| Coding regions |  |  |  |  |  |  |  |  |
| Synonomous | 4 | 0.31 | 4 | 0.17 | 4 | 0.21 | 6 | 0.43 |
| Non synonymous | 9 | 0.69 | 20 | 0.83 | 15 | 0.79 | 7 | 0.5 |
| Amino Acid changes* |  |  |  |  |  |  |  |  |
| Conservative | 3 | 0.33 | 10 | 0.5 | 9 | 0.6 | 1 | 0.14 |
| Non Conservative | 6 | 0.67 | 10 | 0.5 | 6 | 0.4 | 6 | 0.86 |
| **OUTCOME OF SMALL INDELS (<4 Base pairs)** | | | | | | | | |
| Positions |  |  |  |  |  |  |  |  |
| Coding | 1 | 1 | 6 | 1 | 2 | 1 | 0 | 0 |
| Non-coding | 0 | 0 | 0 | 0 | 0 | 0 | 0 | 0 |
| **Outcome of Large Deletions** | | | | | | | | |
| Positions |  |  |  |  |  |  |  |  |
| Coding | 0 |  | 5 | 0.63 | 2 | 1 | 0 | 0 |
| Non coding | 0 |  | 0 |  | 0 | 0 | 0 | 0 |
| Repeat elements | 0 |  | 3 | 0.38 | 0 | 0 | 0 | 0 |

**pMutS, pMutL and pMutS-MutL plasmids were used for overexpression of MutS, MutL or MutS-MutL in 102BW *E.coli* cells respectively; Control represents empty vector control, pBAD18-Kan, carrying 102BW cells.**

***Blosum 62 matrix was used to determine whether mutation is conservative or not**
