## Supplementary table 5 for "Overexpression of MutS impairs DNA mismatch repair and causes cell division defect in *E.coli*"

| **Supplementary table 5 : Spontaneous rate of rifampicin resistance in 102BW *E.coli* cells under conditions mentioned below:** | Rif^R^ Mutation Rate  (X10^9^) | 95% CL  ( X10^9^) |
| --- | --- | --- |
| Control | 2.08 | ±0.85 |
| MutS overexpression | 2.25 | ±0.94 |
| F36A MutS overexpression | 3.0 | ±1.28 |
| MutS-MutL Co-overexpression | 2.04 | ±1.05 |

- **pMutS, pF36AMutS and pMutS-MutL plasmids were used for overexpression of MutS, F36AMutS or MutS-MutL in 102BW E.coli cells. Control is pBAD18-Kan carrying 102BW cells.**
- **For Rif^R^ rate measurement, fluctuation experiment was done with 19 parallel cultures for control and MutS overexpression while for F36A MutS and MutS-MutL overexpression 10 parallel cultures were used.**
- **Mutation rate was determined using web application bz-rates. 95% CL provides 95% confidence limits for the estimated mutation rates.**
