## Supplementary table 6 for "Overexpression of MutS impairs DNA mismatch repair and causes cell division defect in *E.coli*"

| Strain or Plasmid Name  **Supplementary Table 6: Strains and plasmid used in this study** | Genotype | Remarks and/or Reference |
| --- | --- | --- |
| BW27783 | *Δ(araD-araB)567, ΔlacZ4787(::rrnB-3), λ^-^, Δ(araH-araF)570(::FRT), ΔaraEp-532::FRT, φP_cp8_araE535, rph-1, Δ(rhaD-rhaB)568, hsdR514* | ^60^ |
| DY378 | W3110 *ʎcI857 ∆(cro-bioA)* | ^65^ |
| MG1655 | *rph-1* | Lab resource |
| CSH142 | *F^-^,ara-600*, *Δ(gpt-lac)5*, *λ^-^*, *relA1*, *spoT1*, *thiE1* | ^61^ |
| 101MG | MG1655*,E461X lacZ, lacI::Tn10* | measures AT to CG BPS  ; Manjula Reddy (Unpublished) |
| 102MG | MG1655*,E461G lacZ, lacI::Tn10* | measures GC to AT BPS  ; Manjula Reddy (Unpublished) |
| 103MG | MG1655*,E461Q lacZ, lacI::Tn10* | measures GC to CG BPS  ; Manjula Reddy (Unpublished) |
| 104MG | MG1655*,E461A lacZ, lacI::Tn10* | measures GC to TA BPS  ; Manjula Reddy (Unpublished) |
| 105MG | MG1655*,E461V lacZ, lacI::Tn10* | measures AT to TA BPS  ; Manjula Reddy (Unpublished) |
| 106MG | MG1655*,E461K lacZ, lacI::Tn10* | measures AT to GC BPS  ; Manjula Reddy (Unpublished) |
| MR-VRI9 | MG1655*,att ʎ:: P_lac_- FtsZgfp (Ampr),nadA::Tet* | Manjula Reddy (Unpublished) |
| 101BW | BW27783*,E461X lacZ, lacI::Tn10* | measures AT to CG BPS ;This study |
| 102BW | BW27783*,E461G lacZ, lacI::Tn10* | measures GC to AT BPS ; This study |
| 103BW | BW27783*,E461Q lacZ, lacI::Tn10* | measures GC to CG BPS ; This study |
| 104BW | BW27783*,E461A lacZ, lacI::Tn10* | measures GC to TA BPS ; This study |
| 105BW | BW27783*,E461V lacZ, lacI::Tn10* | measures AT to TA BPS ; This study |
| 106BW | BW27783*,E461K lacZ, lacI::Tn10* | measures AT to GC BPS ; This study |
| VRI1 | 102BW*, ∆mutS::amp* | This Study |
| VRI2 | 102BW*, ∆sulA::lacZ* | This Study |
| VRI3 | 102BW*, ∆mutL::amp* | This Study |
| VRI4 | 102BW*,∆mutH::cat* | This Study |
| VRI5 | 102BW*,∆recA::cat* | This Study |
| VRI6 | 102BW*,∆dinB::cat* | This Study |
| VRI7 | 102BW*,att ʎ:: P_lac_- FtsZgfp (Amp^r^),nadA::Tet* | This Study |
| VRI8 | BW27783*, att ʎ:: P_lac_- FtsZgfp (Amp^r^),nadA::Tet* | This Study |
| VRI9 | 102BW*, ∆ruvABC::cat* | This Study |
| VRI10 | 102BW*, dam::Tn9* | This Study |
| NO | MG1655*, E461G lacZ, ∆lacI::FRT,∆cynX::FRT* | This Study |
| IO | MG1655 *E461, INV (E461G laczya) , ∆lacI::FRT,∆cynX::FRT* | This Study |
| GJ1922 | JL2301 *lexA^+^ malB::Tn9* | ^43^ |
| pBAD18Kan | *Ara inducible expression vector,pBR322 origin,Kan^r^* | ^64^ |
| pMutS | *pAra::sRBS:mutS,pBAD18kan* | This Study |
| p-nRBSMutS | *pAra::nRBS:mutS, pBAD18kan* | This Study |
| pF36AMutS | *pAra::sRBS: mutS(F36A),pBAD18kan* | This Study |
| pMutS^N^ | *pAra:: sRBS: mutS(Q15A,Q16A,L18A,R19A),pBAD18kan* | This Study |
| pTB63 | *pFts:: ftsQAZ,pSC01,Tet^r^* | ^49^ |
| pMutS-MutL | *pAra::sRBS:mutS-sRBS:mutL operon,pBAD18kan* | This Study |
| pMutS-MutH | *pAra::sRBS:mutS-sRBS:mutH operon,pBAD18kan* | This Study |
