## Supplementary table 7 for "Overexpression of MutS impairs DNA mismatch repair and causes cell division defect in *E.coli*"

| Primer  **Supplementary Table 7: Primers used in this study** | 5' to 3' sequence | Used for |
| --- | --- | --- |
| EcoMutSF | TACGCTAGCTTAAGAAGGAGATATACCATGAGTGCAATAGAAAATTTCGACGCC | Cloning |
| EcoMutSR | TATGTCGACTTACACCAGGCTCTTCAAGCGATAAATCC | Cloning |
| nRBSMutSF | TACGCTAGCAGGGAACCGGACATAACCCCATGAGTG | Cloning |
| rpoBF | ATCCGTTCCGTTGGCGAAATGGC | Sequencing |
| rpoBR | TCGATACCTGCTTCACCCGGATAC | Sequencing |
| lacZF | ACCGCTACGGCCTGTATGTGGTGGATGAAG | Sequencing |
| lacZR | CATCGCGTGGGCGTATTCGCAAAGGATCAG | Sequencing |
| F36AR | TAAAAGAGCTCATACGCATCACCCATCCGGTAAAACAGCAGGATCTC | Cloning |
| UPMutSF | AACAACGCCTCGTAATGCTCC | Cloning |
| MutSNR | GGGCCGCATATGCGGCCATCATGGGCGTATGGGCGTCG | Cloning |
| MutSNF | TGAAAGCCCAGCATCCCGAGATC | Cloning |
| CynXDiaF | ATCGTTTCTATGAAATGTTGCAGG | Sequencing |
| LacIdiaR | GCACGGGAACCGTTAAAGCTGGAAGC | Sequencing |
| EcoMutLF | TACGCTAGCTTAAGAAGGAGATATACCATGCCAATTCAGGTCTTACC | Cloning |
| EcoMutLR | TACGTCGACTCACTCATCTTTCAGGGCTTTTATCGCC | Cloning |
| EcoMutLF-SalI | TACGTCGACTTAAGAAGGAGATATACCATGCCAATTCAGGTCTTACC | Cloning |
| EcoMutHF | TACGTCGACTTAAGAAGGAGAGCTCATCATGTCCCAACCTCGCCCACTGCTC | Cloning |
| EcoMutHR | TACGTCGACCTACTGGATCAGAAAATGACGGGCCAGTAGTG | Cloning |
| 1585C/a | CGCCTGGGCCGAGTGCGGAGATACAACGTTTGTGCGTAATCTCAGACAG | Recombineering |
| 1586G/t | CCGCCTGGGCCGAGTGCGGAGATATGACGTTTGTGCGTAATCTCAGACA | Recombineering |
| 1595C/a | CGGGTCAGACCGCCTGGGCCGAGTACGGAGATACGACGTTTGTGCGTAA | Recombineering |
| GFP-MutSF | ACTAGCTAGCTTAAGAAGGAGACAAACCATGAGTAAAGGAGAAGAACTTTTCAC | Cloning |
| GFP-MutSR | GGAGCCGCCCGGTTTGTATAGTTCATCCATGCCATGTG | Cloning |
| MutSF | ATGAGTGCAATAGAAAATTTCG | Cloning |
| SulADelF | ATGTACACTTCAGGCTATGCACATCGTTCTTCGTCTGTGTAGGCTGGAGCTGCTTC | Recombineering |
| SulADelR | TTAATGATACAAATTAGAGTGAATTTTTAGCCCGCATATGAATATCCTCCTTAGTTCC | Recombineering |
| MutLDelF | ATGCCAATTCAGGTCTTACCGCCACAACTGGCGAATGTGTAGGCTGGAGCTGCTTC | Recombineering |
| MutLDelR | TCACTCATCTTTCAGGGCTTTTATCGCCGGATGTACATATGAATATCCTCCTTAGTTCC | Recombineering |
| MutHDelF | TCAAGGTATCATGACATGTCCCAACCTCGCCCACTTGTGTAGGCTGGAGCTGCTTC | Recombineering |
| MutHDelR | CTACTGGATCAGAAAATGACGGGCCAGTAGTGCACCATATGAATATCCTCCTTAGTTCC | Recombineering |
| DinBdelF | ATGCGTAAAATCATTCATGTGGATATGGACCATATGAATATCCTCCTTAGTTCC | Recombineering |
| DinBdelF | TCATAATCCCAGCACCAGTTGTCTTTCCATTGTGTAGGCTGGAGCTGCTTC | Recombineering |
| RecAdelF | ATGGCTATCGACGAAAACAAACAGAAAGCGTTGGCTGTGTAGGCTGGAGCTGCTTC | Recombineering |
| RecAdelR | GTTAGTTTCTGCTACGCCTTCGCTATCATCTACCATATGAATATCCTCCTTAGTTCC | Recombineering |
